# Astroglial calcium transfer from endoplasmic reticulum to mitochondria determines synaptic integration

**DOI:** 10.1101/2020.12.08.415620

**Authors:** Roman Serrat, Ana Covelo, Vladimir Kouskoff, Sebastien Delcasso, Andrea Ruiz, Nicolas Chenouard, Carol Stella, Corinne Blancard, Benedicte Salin, Francisca Julio-Kalajzić, Astrid Cannich, Federico Massa, Marjorie Varilh, Severine Deforges, Laurie M. Robin, Diego De Stefani, Arnau Busquets-Garcia, Frederic Gambino, Anna Beyeler, Sandrine Pouvreau, Giovanni Marsicano

## Abstract

Intracellular calcium signaling underlies the astroglial control of synaptic transmission and plasticity. Mitochondria-endoplasmic reticulum contacts (MERCs) are key determinants of calcium dynamics, but their functional impact on astroglial regulation of brain information processing is currently unexplored. We found that the activation of astrocyte mitochondrial-associated CB1 receptors (mtCB1) determines MERCs-dependent intracellular calcium signaling and synaptic integration. The stimulation of mtCB1 receptors promotes calcium transfer from the endoplasmic reticulum to mitochondria through specific mechanisms regulating the activity of the mitochondrial calcium uniporter (MCU) channel. Physiologically, mtCB1-dependent mitochondrial calcium uptake determines the precise dynamics of cytosolic calcium events in astrocytes upon endocannabinoid mobilization. Accordingly, electrophysiological recordings in hippocampal slices showed that genetic exclusion of mtCB1 receptors or specific astroglial MCU inhibition blocks lateral synaptic potentiation, a key example of astrocyte-dependent integration of distant synapses activity. Altogether, these data reveal an unforeseen link between astroglial MERCs and the regulation of brain network functions.

## Introduction

Astrocytes represent a large proportion of brain cells and exert key metabolic, structural, synaptic and protective functions [1–4]. In particular, accumulating evidence supports a bidirectional communication between neurons and astrocytes at the so-called tripartite synapse, formed by pre- and post-synaptic elements surrounded by astroglial processes, thereby modulating information processing and behavior [2–4].

Very little is known about the intracellular astroglial mechanisms required to exert these functions, but it is now clear that calcium dynamics at different subcellular astroglial microdomains are key functional elements of the tripartite synapse [5–7]. Astroglial calcium handling is a highly sophisticated process, which is bidirectionally linked to synaptic activity, and where intracellular organelles such as endoplasmic reticulum (ER) and mitochondria play key active roles [8–11]. For instance, astroglial microdomain signaling has been suggested to involve calcium efflux from mitochondria [8]. Moreover, recent data indicate that the positioning of mitochondria within astrocytic processes and their specific calcium handling properties participate in the regulation of synaptic plasticity [11]. Mitochondria/ER contacts (MERCs) are key player of calcium signaling in many cells [12] and have been recently described in astrocytes [13]. However, their functional involvement in astrocyte calcium signaling and synaptic integration is currently unknown.

One of the most interesting functions of the tripartite synapse is lateral synaptic potentiation (LSP), through which astrocytes, upon neuronal depolarization, are able to enhance synaptic efficacy several tens of micrometers away from the stimulation site [14, 15], thereby contributing to fine-tuned synaptic and circuit integration [16]. LSP requires astroglial intracellular calcium elevations, which critically depend on the endogenous activation of type-1 cannabinoid receptors [15, 17] (CB_1_). Discovered as the main target of synthetic and plant-derived cannabinoid drugs, these G protein-coupled receptors form, together with their endogenous ligands, the so-called endocannabinoid system (ECS), which is a key physiological determinant of synaptic and behavioral functions [18–20]).

Recent evidence indicates that, besides its canonical localization at plasma membranes, functional intracellular CB_1_ receptors can be associated to mitochondria [21–23] (mtCB_1_). The activation of mtCB_1_ receptors decreases metabolic processes in brain cells, negatively affecting memory performance [22], social interactions [23] and likely regulating feeding behavior and neuroprotection [22, 24–26]. In particular, recent anatomical data showed that astrocytes are endowed with a significant proportion of mtCB_1_ receptors that are localized close to synapses [23, 27]. ER-dependent calcium signaling has been proposed as a mechanism, through which astroglial CB_1_ receptors trigger different functions of the tripartite synapse, including LSP [15, 28, 29]. However, the precise astroglial intracellular mechanisms underlying these processes, the potential functional involvement of MERCs and the particular implication of mtCB1 receptors are currently unknown.

Here we asked whether astroglial MERCs-dependent calcium signaling is regulated by the activation of mtCB_1_ receptors and contributes to ECS-dependent synaptic plasticity. The results indicate that stimulation of CB_1_ receptors promotes mitochondrial calcium accumulation in cultured astrocytes and in living animals. In particular, ER-dependent calcium entry into mitochondria is actively promoted by mtCB_1_ receptors and regulates cytosolic calcium dynamics. tt, thereby revealing an unforeseen link between mitochondrial functions, astroglial activity and information processing in the brain.

## Results

### Cannabinoid agonists induce mitochondrial calcium increase in astrocytes *via* mtCB_1_ receptors

Previous reports have shown that CB_1_ receptor activation causes changes in intracellular calcium levels in astrocytes [15, 30, 31]. Thus, we asked whether mtCB_1_ receptors participate in the regulation of astroglial calcium dynamics. To address this issue, we used *CB*_*1*_-KO cultured astrocytes transfected either with full-length CB_1_ or the mutant DN22-CB_1_, which lacks the first 22 aminoacids of the receptor [22, 23]. This mutation strongly reduces the amount of mtCB_1_ receptors in the cells and abolishes the mitochondrial effects of cannabinoid agonists, without altering non-mitochondrial CB_1_ receptor-dependent actions [22, 23]. A semi-quantitative analysis of confocal images of immunofluorescence staining of the transfected proteins revealed that this approach provides a reliable and quantitatively similar expression of either CB_1_ or DN22-CB_1_ in cultured astrocytes (**Figure S1A,B**). We then used super-resolution STED microscopy to analyze both the overlap and the minimal distance between the mitochondrial marker Tom20 (Mitochondrial outer membrane translocase 20 kD) and CB_1_ or DN22-CB_1_ proteins, respectively. The analysis of these STED images revealed that a relatively high proportion of CB_1_ protein either overlapped with (**Figure S1A,C**) or was placed at low distance from Tom20 (**Figure S1D,E**). Conversely, the overlap of DN22-CB_1_ with Tom20 was strongly reduced as compared to CB_1_ (**Figure S1A,C**), and the distance from the mitochondrial marker was significantly higher (**Figure S1D,E**). Thus, the wild-type CB_1_ protein significantly associates to astroglial mitochondria, whereas the DN22-CB_1_ mutant displays much lower mitochondrial association.

To simultaneously record mitochondrial and cytosolic calcium signals, we generated a mitochondrial genetically encoded indicator (Mito-GCaMP6s) and we used it in combination with the cytosolic indicator RCaMP2 [32]. These constructs were co-transfected into *CB*_*1*_-KO astrocytes, together with CB_1_ or DN22-CB_1_ The high affinity cannabinoid agonist WIN 55,212-2 (WIN, 100 nM) increased cytosolic calcium levels in astrocytes expressing the CB_1_ receptor, but not in control (Ctrl) *CB*_*1*_-KO cells (**Figure 1A-C**). This CB_1_ receptor-dependent response was similar in DN22-CB_1_-transfected astrocytes (**Figure 1B-D**), indicating that the deletion of the first 22 aminoacids does not impair the ability of CB_1_ receptors to increase cytosolic calcium levels and that mitochondrial localization of the protein is unlikely to participate in this effect.

**Figure 1.**
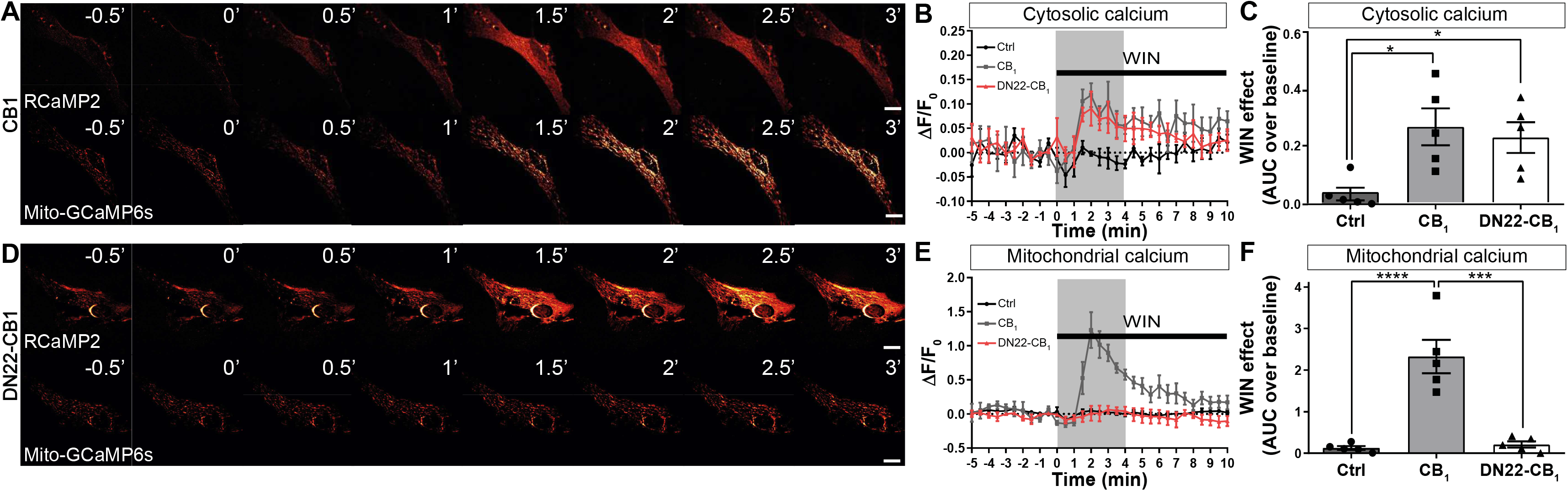
MtCB1 activation increases mitochondrial calcium. **A,D** Representative time-point RCaMP2 (upper panels) and Mito-GCaMP6s (bottom panels) images from CB1 (A) or DN22-CB1 (D) transfected astrocytes. WIN (100 nM) treatment was applied at time 0. Scale bar: 10 μm. **B,E** ΔF/F0 traces from RCaMP2 (B) and Mito-GCaMP6s (E) images. Square gray panels show the quantified periods (4 min) in (C,F). **C,F** WIN effect on calcium (Area under the curve, AUC) from ΔF/F0 traces in (B,E) respectively. Baseline: 1 min before WIN treatment. *p-value<0.05; ***p-value<0.001; ****p-value<0.0001.

Interestingly, the same treatment also induced an increase in mitochondrial calcium levels in astrocytes transfected with wild-type CB_1_, but not in *CB*_*1*_-KO cells (**Figure 1A,E,F**). Strikingly, however, mitochondria from astrocytes expressing the DN22-CB_1_ mutant did not respond to the WIN challenge (**Figure 1D-F**). To rule out possible unspecific effects of our manipulations on the general ability of mitochondria to uptake calcium, the same cells used to study WIN effects were subsequently treated with the well-known astroglial and mitochondrial activator ATP [33] (50 μM). Contrary to WIN, ATP caused a similar mitochondrial calcium response in all three conditions (**Figure S2A**).

Together, these data indicate that mtCB_1_ receptors are dispensable for cannabinoid-induced increase of cytosolic calcium in astrocytes, but they are necessary for the cannabinoid-induced rise of the ion levels in astroglial mitochondria.

### Molecular mechanisms of the mtCB_1_ receptor-dependent mitochondrial calcium increase in astrocytes

Through the activation of inositol triphosphate receptor (IP3R), the endoplasmic reticulum (ER) represents the major source of calcium in astrocytes [34, 35]. Mitochondria physically contact ER at the level of MERCs [36] (mitochondria-ER contacts), allowing the transfer of calcium between these two organelles (**Figure 2A**). MERCs were also present in our *CB*_*1*_-KO astrocyte cultures and their structure was not altered by the expression of CB_1_ or DN22-CB_1_ receptors (**Figure 2B** **and Figure S3A-D**). To investigate the role of ER and MERCs in mtCB_1_ receptor-dependent mitochondrial calcium increase, we first treated CB_1_-expressing astrocytes with the blocker of the sarco/endoplasmic reticulum calcium-ATPase (SERCA) pump Thapsigargin (TG, 1 μM), which empties the ER from calcium (**Figure 2A**). TG treatment fully occluded the calcium responses to WIN either in the cytosol (**Figure 2C** **and Figure S4A**) or mitochondria (**Figure 2D** **and Figure S4B**). Thus, the ER mediates both cytosolic and mitochondrial cannabinoid-induced calcium increases.

**Figure 2.**
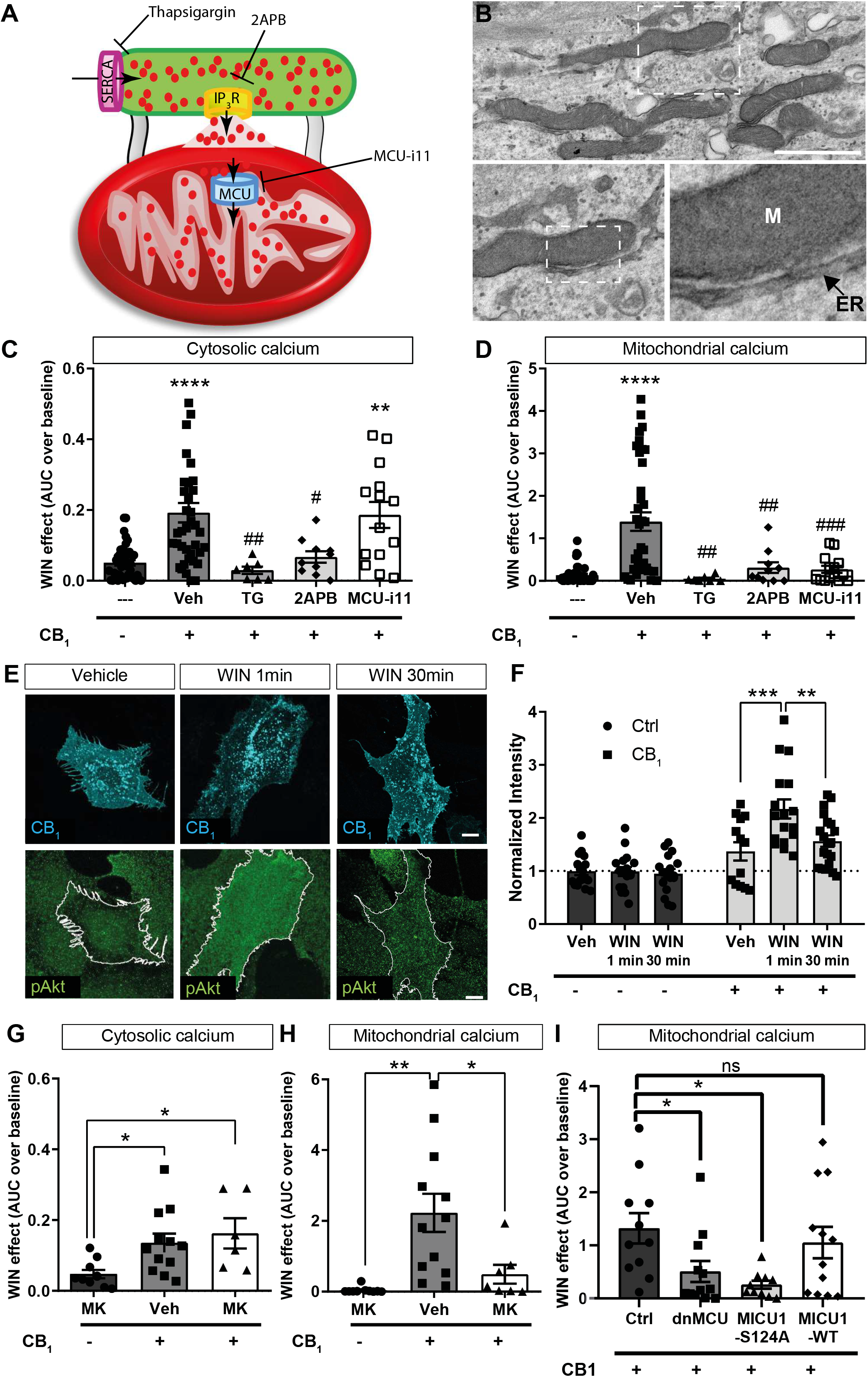
ER/IP3R/MCU and AKT pathways are necessary for CB1-dependent mitochondrial calcium increase. **A** Scheme showing ER (in green) -mitochondria (in red) calcium transfer and the different drugs used in Figure 2 to study the role of this pathway. **B** Representative electron microscopy image from CB1-KO astrocytes. M: mitochondria. ER: endoplasmic reticulum. Scale bar: 0,5 μm. **C,D** WIN effect on cytosolic (D) and mitochondrial (D) calcium (Area under the curve, AUC) in negative control (CB_1_−), Vehicle control (Veh), thapsigargin (TG), 2APB and MCU inhibitor 11 (MCU-i11) conditions. Baseline: 1 min before WIN treatment. ΔF/F0 traces are shown in Figure S4A,B respectively. **E** Representative images from CB1 transfected astrocytes in Vehicle, WIN-1 min and WIN-30 min conditions. In cyan, CB1; in green, p-AKT (T296). Scale bar: 10 μm. **F** Normalized total intensity from cells show in (E) in control CB_1_− (Ctrl) and CB_1_+ transfected conditions. **G,H** WIN effect on cytosolic (G) and mitochondrial (H) calcium (AUC) in negative control CB_1_−, Vehicle control (Veh) and AKT-inhibitor (MK) conditions. Baseline: 1 min before WIN treatment. ΔF/F0 traces are shown in Figure S4C,D respectively. **I** WIN effect on mitochondrial calcium (AUC) in CB_1_+ trasfected cells co-trasnfected with dnMCU, MICU1-S124A or MICU1-WT. Baseline: 1 min before WIN treatment. ΔF/F0 traces are shown in Figure S4E. *p-value<0.05; **p-value<0.01; ***p-value<0.001; ****p-value<0.0001. #p-value<0.05; ##p-value<0.01; ###p-value<0.001 in comparison to Vehicle (CB_1_+) condition.

IP3R are the main players of metabotropic control of cytosolic calcium in astrocytes and, interestingly, they are present in MERCs [34, 35, 37]. To determine whether IP3R are involved in cannabinoid-mediated increase in cytosolic and/or mitochondrial calcium, we applied the IP3R antagonist 2-Aminoethoxydiphenyl borate (2APB, 100 μM, **Figure 2A**) to CB_1_-expressing astrocytes. 2APB drastically reduced both CB_1_ receptor-dependent cytosolic and mitochondrial calcium responses (**Figure 2C**,**D** **and Figure S4A,B**), indicating that both effects require the release of calcium from the ER through IP3R.

The main channel for the entry of calcium into the mitochondrial matrix is the mitochondrial calcium uniporter [9] (MCU, **Figure 2A**). The application of the MCU-inhibitor 11 [38] (MCU-i11, 10 μM) to CB_1_-expressing astrocytes did not alter the amplitude of the increase in bulk cytosolic calcium induced by WIN (**Figure 2C** **and Figure S4A**). However, this treatment strongly reduced WIN-induced increase in mitochondrial calcium levels (**Figure 2D** **and Figure S4B**), indicating that MCU activity is specifically required for this mtCB_1_ receptor-dependent effect.

MtCB_1_ receptors have been recently proposed to activate protein kinase B/AKT [26] (hereafter called AKT). Interestingly, recent data indicate that AKT stimulation increases the entrance of calcium in the mitochondrial matrix by phosphorylating the mitochondrial calcium uptake protein 1 (MICU1), an important MCU modulator [39]. Specific immunocytochemistry approaches revealed that WIN treatment of CB_1_-expressing astrocytes caused a fast and transient increase in AKT phosphorylation levels (p-AKT; **Figure 2E,F**), indicating that activation of CB_1_ receptors triggers the AKT pathway in cultured astrocytes. The application of the AKT blocker MK-2206 (10 μM) did not impair CB_1_ receptor-dependent cytosolic calcium increase (**Figure 2G** **and Figure S4C**), but it reduced the WIN effect on mitochondria (**Figure 2H** **and Figure S4D)**. To further investigate the potential involvement of MCU and the AKT/MICU1 signalling, genetically blockade of this pathways was investigated. Astrocytes were co-transfected with a dominant negative mutant version of MCU [40] (dnMCU) or the non-phosphorylatable form MICU1-S124A [39]. The expression of either dnMCU or MICU1-S124A effectively blunted WIN effect on mitochondrial calcium (**Figure 2I** **and Figure S4E**).

Altogether, these data indicate that the activation of IP3R through non-mitochondrial CB_1_ receptors is sufficient for cannabinoid-induced cytosolic calcium increase in astrocytes. Conversely, in addition to stimulation of IP3R, mtCB_1_ receptors, AKT signaling, MICU1 phosphorylation and MCU activity are required for cannabinoid-induced mitochondrial calcium uptake.

### CB_1_ receptor activation increases mitochondrial calcium in astrocytes *in vivo*

Many differences exist between cultured astrocytes and the same cells in living animals [41]. To study if CB_1_ receptor-dependent mitochondrial calcium responses occur also *in vivo,* we generated an AAV virus expressing the sensor Mito-GCaMP6s under the promoter of the human glial fibrillary acidic protein [42] (hGFAP, to generate AAV-GFAP-Mito-GCAMP6s). The injection of this virus into the somatosensory cortex or the hippocampus resulted in a specific expression of the calcium sensor in astrocytes (**Figure 3A**,**B** **and Figure S5A**). Four weeks after surgery, anesthetized head-fixed wild-type and *CB*_*1*_-KO mice carrying astroglial Mito-GCaMP6s expression in the somatosensory cortex were treated with the plant-derived cannabinoid Δ^9^-tetrahydrocannabinol (THC, 10 mg/kg) and fluorescence levels were analyzed by two-photon microscopy (**Figure 3C,D**). Time course image analysis (**Figure 3E**) revealed that the THC treatment progressively increased the occurrence of mitochondrial calcium events in wild-type, but not in *CB*_*1*_-KO mice (**Figure 3F**). In particular, whereas a non-significant trend effect of THC was already observed during the first 15 min post-injection, the increase in the number of calcium events became evident at min 15 to 30, with no changes in *CB*_*1*_-KO mice (**Figure 3G,H**). These data indicate that THC increases astroglial mitochondrial calcium levels *in vivo*.

**Figure 3.**
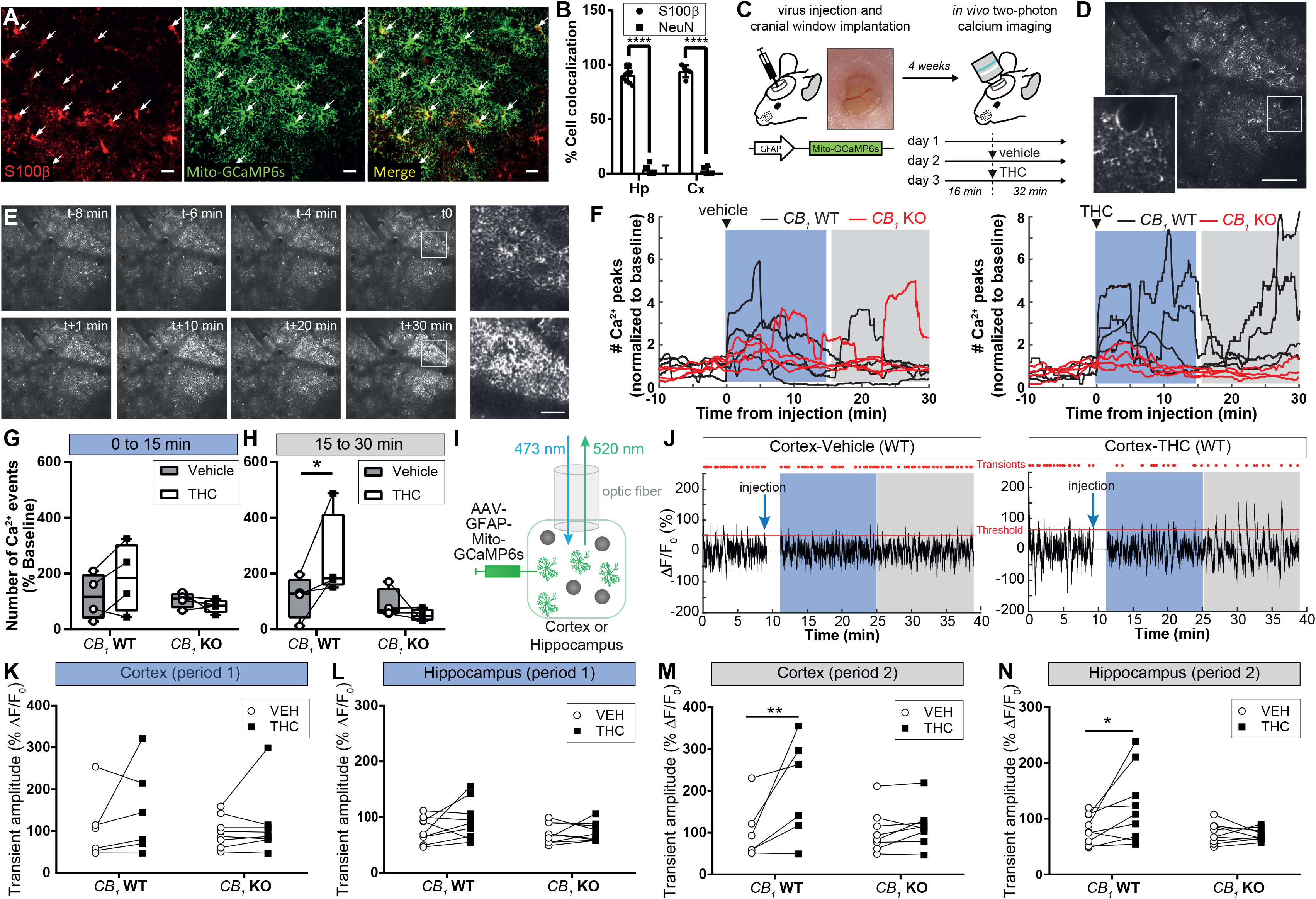
CB1 activation induces mitochondrial calcium increase *in vivo*. **A** Immunohistochemistry representative images from mouse hippocampi injected with AAV-GFAP-Mito-GCaMP6s (in green). Astrocyte marker S100β (in red). Scale bar: 20 μm. **B** Percentage of cell colocalization of Mito-GCaMP6s and S100β (astrocytic marker) or NeuN (neuronal marker) in Hippocampi (Hp) and Cortex (Cx). **C** Experimental two-photon imaging approach for *in vivo* recording of mitochondrial calcium dynamics in mouse somatosensory cortex. **D** Representative two-photon image. Squared image shows mitochondrial particles (yellow arrow heads). Scale bar: 100 μm. **E** Representative time-point two-photon images of the somatosensory cortex from *CB*_*1*_ WT mouse injected with AAV-GFAP-Mito-GCaMP6s. THC (10 mg/kg) treatment was applied at time 0. On the right magnified images from time 0 and time 30 min (white squares). Scale bar: 30 μm. **F** Normalized calcium peak traces from *CB*_*1*_ WT and *CB*_*1*_ KO mice injected with Vehicle or THC. Blue and gray rectangles show the time window analyzed in G and H. **G,H** Average percentage of calcium events (normalized to day 1) between 0 to 15 min (G) or 15 to 30 min (H) in *CB*_*1*_ WT and *CB*_*1*_ KO mice treated with Vehicle or THC. **I** Scheme showing experimental fiber photometry imaging approach. **J** Representative traces from the cortex of a *CB*_*1*_ WT mouse injected with AAV-GFAP-Mito-GCaMP6s. Vehicle-injected (left panel) or THC-injected (10 mg/kg, right panel) mouse traces. Red dots correspond to detected transients above threshold (median+2*MAD). Blue and gray rectangles show the time window of the analyzed “period 1” and “period 2” respectively. Note that the first minute before and after injection was removed to exclude mouse-handling/injection effects. **K,M** Cortical or **L,N** hippocampal amplitude of transients in *CB*_*1*_ WT and *CB*_*1*_ KO mice treated with Vehicle (VEH) or THC during the 2 time periods highlighted in (J). *p-value<0.05; **p-value<0.01; ****p-value<0.0001.

To investigate the effect of cannabinoid treatment in multiple brain regions simultaneously and in freely moving animals, we performed fiber photometry recordings in the hippocampus and somatosensory cortex of mice carrying astroglial Mito-GCaMP6s expression (**Figure 3A**,**B** **and Figure S5A**), treated with vehicle or THC (**Figure 3I,J**). The amplitude (Am), duration (Du) and frequency (Fq) of recorded calcium events were analyzed in 2 equal time periods after vehicle or THC injection (10 mg/kg i.p.; see Methods). No significant differences were observed during the first 15 min of recording (**Figure 3K**,**L** **and Figure S5B,C**). However, during the second half of recording, THC significantly increased the amplitudes of astroglial mitochondrial calcium signals in both hippocampus and cortex of *CB_1_-*WT mice but not of *CB*_*1*_-KO littermates (**Figure 3M,N**). Despite some non-significant trend, frequency and duration of transients were not altered by the drug (**Figure S5D,E**). Thus, systemic THC administration increases astroglial mitochondrial calcium levels in both hippocampus and somatosensory cortex of freely moving animals.

### Physiological role of mtCB1 receptors in astrocyte-dependent synaptic integration

Neuronal depolarization induces the mobilization of endocannabinoids leading to retrograde suppression of neurotransmitter release [19, 20]. However, these endocannabinoids can also activate CB_1_ receptors in neighboring astrocytes and, by increasing intracellular calcium activity, induce distal glutamate release and lateral synaptic potentiation [15-17, 31] (LSP). To investigate the potential role of mtCB_1_ receptors in this phenomenon, we performed simultaneous whole-cell electrophysiological recordings of two CA1 pyramidal neurons in acute hippocampal slices (> 60 μm apart, termed “proximal” and “distal” neurons). Upon minimal stimulation of the Schaffer collaterals [15], the activity of putative single synapses onto the distal neuron was recorded. After neuronal depolarization (5 s at 0 mV) of the proximal neuron to induce endocannabinoid mobilization [15], we analyzed changes in synaptic parameters recorded in the distal neuron (**Figure 4A**). As expected [15], neuronal depolarization induced an increase in the success rate in 9 out of 19 distal neurons derived from wild-type animals (47.4%; **Figure 4B-E**), without changes in the amplitude of evoked excitatory post-synaptic potentials (EPSCs; **Figure 4B** **and Figure S6A**), thereby indicating the occurrence of LSP. We then used a recently generated knock-in mouse mutant line, where the DN22-CB_1_ sequence replaces the wild-type CB_1_ gene [43] (DN22-CB_1_-KI mice). These mutants lack the anatomical association of the CB_1_ protein with mitochondria, and do not display typical *in vivo* and *ex vivo* mitochondrial-related effects of cannabinoids [43] (*e.g.* decrease of cellular respiration and amnesic effects). However, the mutant mice maintain other functions of CB_1_ receptors, such as the pharmacological activation of G protein signaling or the physiological regulation of hippocampal inhibitory neurotransmission [43]. Interestingly, basal synaptic parameters were not altered in hippocampal slices obtained from DN22-CB_1_-KI mice (**Figure S6B,C**), whereas LSP was virtually absent (**Figure 4C-E** **and Figure S6A**), indicating that mtCB_1_ receptors are necessary for this synaptic phenomenon.

**Fig 4.**
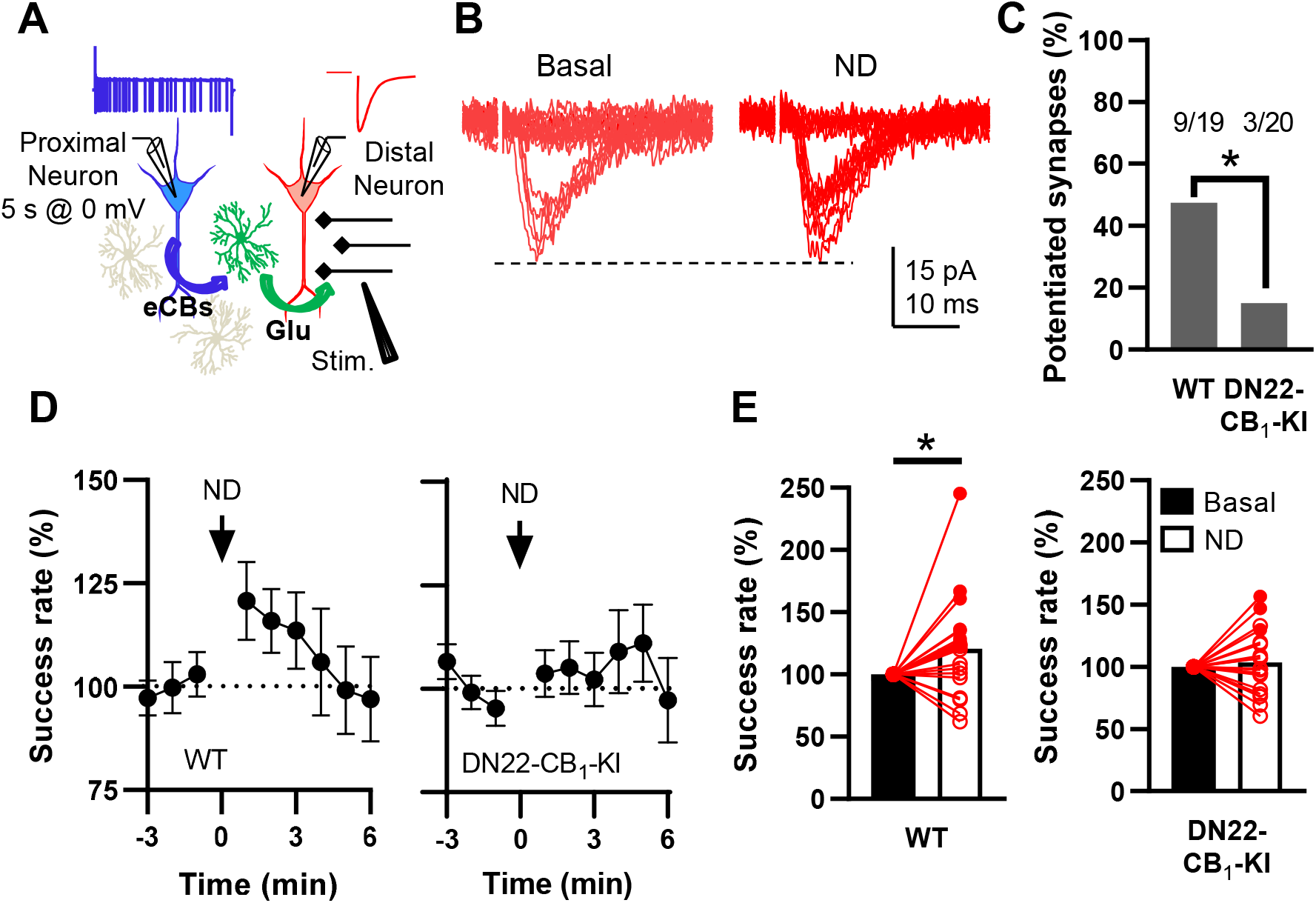
MtCB_1_ participates in lateral synaptic potentiation. **a** Scheme depicting the experimental approach. eCBs: endocannabinoids. Glu: glutamate. **b** Representative traces obtained before (Basal) and after neuronal depolarization (ND). **c** Percentage of potentiated synapses recorded in WT and DN22-CB_1_-KI hippocampal slices. **d** Time course of normalized success rate in WT and DN22-CB_1_-KI conditions. Arrow indicates neuronal depolarization (ND). **e** Normalized success rate before (Basal) and after neuronal depolarization (ND) time in WT and DN22-CB_1_-KI conditions. Red and white circles represent potentiated and non-potentiated synapses, respectively. *p-value<0.05.

### Mitochondrial calcium uptake in astrocytes is essential for mtCB1 receptor-dependent lateral synaptic potentiation

The results obtained so far indicate that mtCB_1_ receptors are necessary for physiological endocannabinoid-dependent LSP in the hippocampus and for the pharmacological cannabinoid-induced and MCU-dependent increase of mitochondrial calcium in astrocytes. Thus, we asked whether mitochondrial calcium regulation might participate in the astroglial signaling underlying LSP [15]. Upon LSP-inducing neuronal depolarization, cytosolic calcium dynamics were recorded in hippocampal slices expressing the GCaMP6f indicator in astrocytes (**Figure 5A** **and Figure S7A,B**). Neuronal depolarization effectively induced an increase in the amplitude, frequency, spreading and duration of large astroglial calcium events (>40 μm^2^ of spreading coefficient, see Methods, **Figure 5B-E**), without affecting smaller ones (**Figure S7C-F**). Interestingly, whereas the application of MCU-i11 did not affect the depolarization-induced increase of astroglial calcium events amplitude (**Figure 5B** **and Figure S7B**), it abolished the effects on frequency, spreading and duration (**Figure 5C-E**). These data indicate that mitochondrial calcium handling specifically impacts the dynamic changes of astroglial calcium signals accompanying neuronal depolarization-induced LSP.

**Fig 5.**
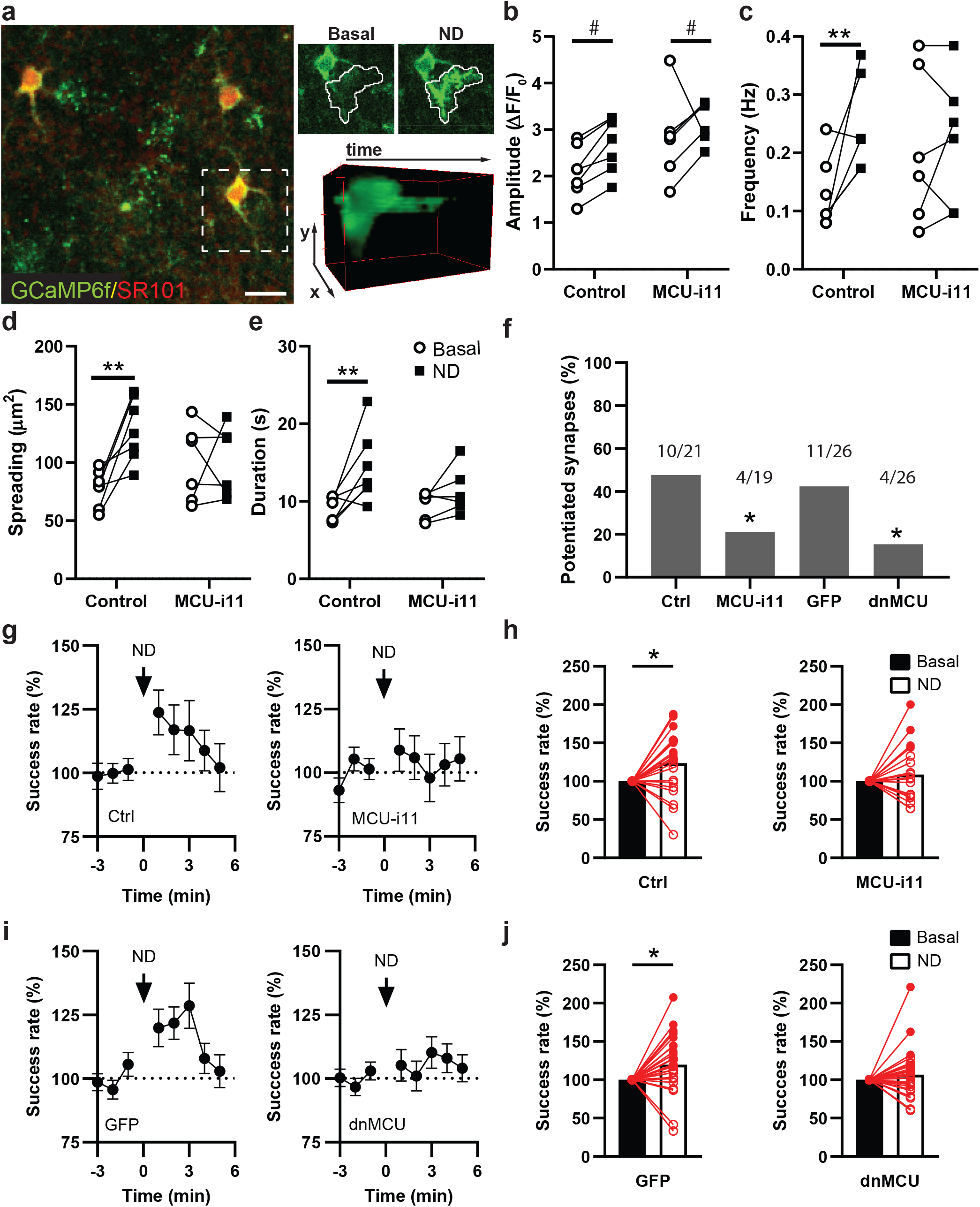
MCU blockade disturbs astrocyte-neuron signaling. **a** Left panel: Representative confocal image from mouse hippocampi injected with AAV-GFAP-GCaMP6f (in green). Astrocyte marker SR101 (in red). Scale bar: 20 μm. Right panels: Localization (white line) of a calcium event before or after ND and 3D representation of the event in x, y and t (see Methods). Graphs obtained from calcium confocal imaging in hippocampal slices (a) showing **b** mean amplitude, **c** frequency, **d** mean spreading coefficient and **e** mean duration of large calcium events (>40 μm^2^) before (Control) or 20 min after MCU-i11 treatment. ND: neuronal depolarization. **f** Percentage of potentiated synapses recorded in different experimental conditions. **g** Time course of normalized success rate in control (Ctrl) and MCU-i11 conditions. Arrow indicates neuronal depolarization (ND). **h** Normalized success rate before (Basal) and after neuronal depolarization (ND) time in control (Ctrl) and MCU-i11 conditions shown in (g). Red and white circles represent potentiated and non-potentiated synapses, respectively. **i,j** Like (g) and (h) but from mice hippocampal slices injected with GFP or with dnMCU. *p-value<0.05; **p-value<0.01. #p-value<0.05 general “stimulation” effect on amplitude.

To further investigate whether this function of mitochondria participates in mtCB_1_ receptor-dependent LSP, we directly addressed the role of MCU in this phenomenon. MCU-i11 strongly reduced the number of neurons undergoing depolarization-induced LSP (**Figure 5F**), abolishing the increase of success rate onto the distal neuron (**Figure 5G,H**), without altering the amplitude of EPSCs (**Figure S8A**). These results suggest that MCU activity is required for LSP in the hippocampus. However, control experiments revealed that MCU-i11 did not alter the amplitude of basal EPSCs (**Figure S8B**), but it slightly increased basal success rate (**Figure S8C**), suggesting that the blockade of LSP might be confounded by an occlusion effect of the drug. Moreover, MCU is expressed in both astrocytes and neurons [44], leaving the possibility that its blockade impairs LSP through other mechanisms than a direct mtCB_1_ receptor-dependent regulation of astroglial mitochondrial calcium. To control for these potential confounders, we adopted a genetic approach, and we generated an AAV carrying the expression of dnMCU in astrocytes *in vivo* (AAV-GFAP-dnMCU; **Figure S8D**). The expression of dnMCU in hippocampal astrocytes did not alter basal EPSC amplitude, or the success rate (**Figure S8B,C**). However, strikingly, the astroglial expression of dnMCU blocked LSP (**Figure 5F,I,J** and **Figure S8A**), thereby confirming the functional importance of MCU and excluding potential unspecific and/or non astroglial effects of pharmacological inhibition.

Altogether, these data show that mitochondrial calcium uptake by MCU controls both endocannabinoid-dependent cytosolic calcium dynamics and mtCB_1_-dependent LSP.

## Discussion

This study reveals an unforeseen mechanism linking mitochondrial and MERCs calcium dynamics in astrocytes and synaptic integration in hippocampal circuits. In particular, activation of mitochondrial-associated cannabinoid receptors (mtCB_1_) in astrocytes promotes ER- and IP3R-dependent mitochondrial calcium entry, requiring AKT signaling, MICU1 phosphorylation and MCU activity. Moreover, endogenous physiological activation of mtCB_1_ receptors by neuronal depolarization regulates astroglial cellular calcium dynamics, and thereby mediates lateral potentiation of excitatory synaptic transmission. Therefore, mtCB_1_ receptor regulation of MERCs functions is an important element of astroglial control of synaptic integration.

Besides other roles, mitochondria are known to uptake calcium, contributing to the shaping of cytosolic calcium dynamics and, thereby, to the regulation of several important cellular processes [45–48]. Mitochondrial calcium uptake is generally considered as a passive mechanism regulated only by cytosolic calcium levels [49], although recent evidence suggested that this process might be actively modulated [39, 50, 51]. Our data clearly show that mtCB1 receptors actively promote mitochondrial calcium intake in astrocytes, independently of cytosolic levels. In addition to CB_1_, other metabotropic receptors have been proposed to directly regulate mitochondrial functions [52–55]. For instance, the stimulation of P2Y2 purinergic receptors in *in vitro* isolated mitochondria can increase mitochondrial calcium levels [53]. Our data show that the effects of cannabinoids on astroglial mitochondrial calcium levels are present both in cultured cells and *in vivo,* demonstrating that this function of metabotropic receptors is relevant to living and behaving animals. The novel possibility to record mitochondrial calcium *in vivo* is based on viral approaches, which is inevitably accompanied with a certain degree of gliosis. We cannot, therefore, exclude that reactive astrocytes are the ones interested by our recordings. Nevertheless, our data state the principle that THC can increase mitochondrial calcium levels *via* CB1 in astrocytes of living and freely moving animals.

By showing that mtCB_1_ receptors are necessary for cannabinoid-induced mitochondrial but not cytosolic calcium increases, our data reinforce the idea that specific signaling cascades actively modulate calcium dynamics in these organelles. In particular, distinct subcellular pools of CB_1_ receptors differentially participate in the modulation of subcellular calcium dynamics. The observation that inhibition of AKT or the expression of a non-phosphorylatable form of MICU1 reduces mitochondrial but not cytosolic cannabinoid-mediated increases of calcium further supports this idea. Altogether, these data reveal that mtCB_1_ receptors represent an active regulator of dynamic mitochondrial calcium uptake in astrocytes, through a molecular cascade involving IP3R, AKT, MICU1 and MCU.

CB_1_ receptor-dependent cytosolic calcium increase in astrocytes mainly relies on the release of calcium from the ER (present results and Refs. [15, 31]). Our data indicate that ER calcium release is also needed for mtCB_1_ receptor-dependent mitochondrial calcium increase. In our EM analysis of cultured astrocytes, almost 100% of mitochondria are located less than 100 nm from ER, a distance that permits calcium exchanges between the 2 organelles and defines MERCs [36]. In addition, previous EM studies of hippocampal slices have also revealed (i) that astroglial mtCB_1_ receptors are in the vicinity of synapses [27] (less than 1 μm away) and (ii) that MERCs are present in the astrocytic processes [13]. Hence, our data support the notion that mtCB_1_ receptors are placed in the ideal position to regulate MERCs-dependent calcium transfer and thereby modulate astrocytic calcium signaling.

Amplification sites for calcium waves in astrocytes are characterized by the presence of ER calcium signaling proteins (*e.g.* calreticulin and IP3R), and of mitochondria [10], suggesting that MERCs calcium transfer could be needed for the propagation of cytosolic calcium events. Interestingly, abolition of cannabinoid-induced mitochondrial calcium entry by blocking MCU, removing mtCB1 or inhibiting AKT does not affect the amplitude of the CB1-dependent simultaneous cytosolic calcium increase in cultured astrocytes. This could be due to technical reasons. For instance, the indicator used to detect cytosolic calcium RCaMP2 has a low dynamic range, which could prevent the detection of small changes in amplitude [32]. However, the use of the larger dynamic range indicator GCaMP6f to detect cytosolic calcium in hippocampal slices also failed to reveal any difference in the amplitude of endocannabinoid-induced cytosolic calcium events in similar conditions. Thus, lack of effect of the blockade of mitochondrial calcium uptake on the amplitude of cytosolic calcium transient seems to reflect physiological mechanisms rather that technical limits of the experiments. Indeed, it has been shown that mitochondria might act as poor dynamic buffer of bulk cytosolic calcium under physiological conditions especially when other strong extrusion systems, such as plasma membrane and ER calcium pumps, are available, whereas their activity still regulates cytosolic calcium levels in specific microdomains [56, 57]. Further studies on the astrocytic calcium machinery would be needed to dissect the role of mitochondria in the regulation of the amplitude of astrocytic calcium transient under physiological conditions.

Conversely, our data obtained in hippocampal slices show that MCU inhibition affects the spatial and temporal spreading of astroglial cytosolic calcium events after activation of endocannabinoid signaling by neuronal depolarization. This indicates that mtCB1-dependent regulation of MERCs functions plays a key physiological role in the dynamic properties of calcium signaling. Mitochondria, however, have been proposed to modulate calcium waves propagation in the opposite way, with impairment of mitochondrial function increasing the spreading and duration of calcium signals [58–60]. As these studies used strong pharmacological stimulation of ER calcium release whereas our data are based on physiological endogenous stimulations, this apparent discrepancy could originate from differences in methodological approaches. However, it is also possible that the mtCB_1_ receptor-dependent mitochondrial calcium increase exerts a specific signaling function. For instance, it might prevent the negative feedback of IP3R following changes in local calcium concentrations, which has been proposed to regulate cytosolic calcium spreading [61]. Altogether, our results suggest that ER/mitochondria calcium transfer induced by activation of mtCB_1_ receptors is important for the propagation, but not the amplitude, of cytosolic calcium waves in astrocytes.

Over the course of the last 20 years, numerous studies have led to the conclusion that neuronal network functions result from the coordinated activity of astrocytes and neurons [16]. It is well known that astroglial intracellular calcium elevations induced by synaptic transmission cause in turn a modulation of neuronal activity and synaptic plasticity [5, 62–64]. One example of this phenomenon is the astroglial CB_1_ receptor-induced LSP of excitatory synapses [15, 16]. Our data show that the control of calcium dynamics by the activation of mtCB_1_ receptors is required to express LSP. By allowing the coordinated activity of neurons that are not necessarily synaptically connected, LSP likely represents a mechanism to expand the potential computational impact of neuronal networks [16]. Therefore, astroglial mtCB_1_ receptor-dependent control of MERCs functions actively participates in synaptic plasticity and likely contributes regulating complex information processing in the brain.

Overall, our findings unravel the importance of astroglial mitochondria and MERCs in shaping neuronal network activity and uncover a novel mechanism of action of CB_1_ receptor signaling in the brain.

## Acknowledgments

The microscopy was done in the Bordeaux Imaging Center a service unit of the CNRS-INSERM and Bordeaux University, member of the national infrastructure France BioImaging supported by the French National Research Agency (ANR-10-INBS-04). The help of Christel Poujol, Sébastien Marais and Fabrice Cordelières is acknowledged. We also thank Delphine Gonzales, Nathalie Aubailly and all the personnel of the Animal Facilities of the NeuroCentre Magendie for mouse care. We thank the Biochemistry Platform of Bordeaux NeuroCampus for help. We also thank all the members of Marsicano’s lab for useful discussions and for their invaluable support. We thank Manuel Rojo for support in electron microscopy. We thank Haruhiko Bito for providing Rcamp2 plasmid. We thank Paolo Pinton for providing MICU1 plasmids.We thank Guillaume Ferreira for support, and Jean-Luc Morel and Alfonso Araque for critically reading the manuscript. This work was funded by: INSERM, European Research Council (Endofood, ERC–2010–StG–260515 and CannaPreg, ERC-2014-PoC-640923, MiCaBra, ERC-2017-AdG-786467), Fondation pour la Recherche Medicale (FRM, DRM20101220445), the Human Frontiers Science Program, Region Nouvelle Aquitaine, Agence Nationale de la Recherche (ANR, NeuroNutriSens ANR-13-BSV4-0006 and ORUPS ANR-16-CE37-0010-01) and BRAIN ANR-10-LABX-0043, to GM; Labex Brain (CANNACALC), to SP and F M, and Orups ANR-16-CE37-0010, to R.S; Eu-Fp7 (FP7-PEOPLE-2013-IEF-623638) and MINECO from AEI (RYC-2017-21776), to A.B-G;

## Author contributions

RS and CS performed and analyzed live cell-imaging recordings. ACo performed and analyzed electrophysiological experiments. AR performed and AR and RS analyzed STED-microscopy experiments. CB, BS performed and RS analyzed electron microscopy experiments. VK, NC and FG performed and analyzed two-photon recordings. RS, SDel AB-G and AB performed and analyzed fiber photometry experiments. ACo and SP performed and RS and SP analyzed cytosolic calcium experiments. FJ-K and MV performed immunohistochemistry experiments. ACa contributed to experiments using viral vectors. SDef contributed to plasmid cloning. RS, ACo, GM and SP wrote the manuscript. RS, GM and SP conceived and supervised the whole project and supervised the writing of the manuscript. All authors edited and approved the manuscript.

## Methods

### Mice

All experiments were performed in accordance with the Guide for the Care and Use of Laboratory Animals (National Research Council Committee (2011): Guide for the Care and Use of Laboratory Animals, 8th ed. Washington, DC: The National Academic Press.) and the European Communities Council Directive of September 22th 2010 (2010/63/EU, 74). Experiments were approved by the local ethical committee of the University of Bordeaux (approval number 501350-A) and the French Ministry of Agriculture and Forestry (authorization number 3306369). *CB*_*1*_ WT and *CB*_*1*_ KO male littermate mice were used for two-photon experiments and fiber photometry experiments. Cages were enriched and mice were maintained under standard conditions (food and water ad libitum; 12 h–12 h light–dark cycle. Fiber photometry experiments were done during dark cycle (light off at 8:00a.m.). The rest of the experiments were performed between 9:00 and 19:00 (light on at 7:00). For primary cell cultures, pups were obtained from homozygote *CB*_*1*_ KO pairs. No method of randomization to assign experimental groups was performed and the number of mice in each experimental group was similar. No statistical methods were used to predetermine sample size.

### Drugs

THC was obtained from THC Pharm GmbH (Frankfurt, Germany). WIN55-212-2 (WIN), thapsigargin and ATP disodium salt-hydrate (ATP) were purchased from Sigma-Aldrich (St-Louis, USA); 2APB from Tocris (Bristol, UK); MCU-i11 from ChemDiv (San Diego, USA) and MK-2206 from Enzo Life Sciences (Farmingdale, USA). For in vitro experiments, WIN, 2APB, thapsigargin, MCU-i11 [38] and MK-2206 were dissolved in DMSO; ATP was dissolved in water. DMSO was never higher than 0.001%. For in vivo administration, THC was prepared freshly before the experiments and was dissolved in a mixture of 5% ethanol, 4% cremophor and saline. Corresponding vehicle solutions were used in control experiments. Doses and concentrations of the different drugs were chosen on the basis of previous published data or preliminary experiments.

### Plasmids

The plasmid coding for Mito-GCaMP6s was obtained using standard cloning techniques. Briefly, GcaMP6s (Addgene) starting codon was mutated and the resulting sequence was subcloned into a plasmid containing the mitochondria targeting sequence (MTS) derived from the subunit VIII of human cytochrome C oxidase [65, 66]. RCaMP2 was a generous gift from Prof. H. Bito [32]. The N-terminal deletion of the first 22 aminoacids in the mouse CB_1_ receptor coding sequence to obtain DN22-CB_1_ mutant was achieved as previously described in [22]. The plasmid encoding for the dominant negative version of MCU in pcDNA3.1 plasmid (MCU-D260N-E263Q, dnMCU [40]) was a generous gift from Dr. D. De Stefani (University of Padova, Italy). The plasmids encoding for MICU1-WT and MICU1-S124A [39] was a generous gift from Prof. Paolo Pinton (University of Ferrara, Italy)

### Preparation of adeno-associated viruses (AAV)

AAV-GFAP-Mito-GCaMP6s and AAV-GFAP-dnMCU-IRES-mRuby viral constructs were subcloned using standard molecular cloning techniques previously used in the lab [22]. The resulting vector was transfected with PEI into HEK293 cells together with the AAV8-serotype-packaging plasmids [42], the viruses were then purified by lodixanol desnity gradient and titred as previously described [67]. Virus titres were between 10^10^ and 10^11^ genomic copies per ml for all batches of virus used in the study.

### Cell culture and transfection

Primary cortical astroglial cultures were prepared from *CB*_*1*_ KO P0–P1 mice. Briefly, brains were extracted in PBS containing 0.6% glucose and 0.5% bovine serum albumin (BSA), the cortex were dissected. To dissociate cells 0.25% trypsin (Invitrogen, France) and DNase (Roche, Basel, Switzerland) were used in HBSS 1x. Cells were seeded into 75 cm^2^ flask and after 2 weeks transferred to glass coverslips for live imaging and immunocytochemistry analysis. Astrocytes were kept in MEM (Gibco, France) supplemented with 10% fetal bovine serum (FBS), 2 mM l-glutamine, 120 μg*ml−1 penicillin, 200 μg*ml−1 streptomycin, 1 mM pyruvate, 0.6% Glucose, Fungiozione (Invitrogen, France), and were maintained at 37 °C in the presence of 5% CO2. Transfection was carried out at 1-2 days after coverslip seeding using a standard calcium phosphate transfection protocol, with a 1:1:4 DNA ratio of plasmids (RCaMP2: Mito-GCaMP6s: and empty vector, CB_1_ or DN22-CB_1_, respectively). For the experiments using the dnMCU or MICU1 under CMV promoter, this was used substituting RCaMP2. Astrocytes were cultured for 2 days after transfection before the recordings. Parallel immunocytochemistry showed a high degree of co-transfection of the cells expressing Mito-GCaMP6s and RCaMP2 with the non-tagged CB_1_ vector (not shown). For anatomical studies myc tagged versions of CB_1_ and DN22-CB_1_ were used. For electron microscopy experiments, cells were infected at day 0 using AAV vectors encoding for CB_1_, DN22-CB_1_ or empty vector. 7 days after the infection were fixed as described below.

### Immunocytochemistry

Astrocytes were fixed in 4% formaldehyde dissolved in PBS (0.1 M, pH 7.4). Cells were pre-incubated in a blocking solution (10% normal goat serum, 0.1% triton X-100 and 0.2 M glycine in PBS) for 1 h and then incubated with primary antibody in the same blocking solution for 2 h. Primary antibodies were rabbit anti-TOM20 (1:500; Santa Cruz, sc-11415), mouse anti-myc (1:500; Roche, 11667149001), rabbit anti p-AKT (T296) (1:100; Cell Signalling Technology, 9614) or guinea pig anti-CB_1_ (1:300; Abyntek, CB_1_-gp-AF530). Secondary fluorescent antibodies anti-mouse Alexa488 or anti-rabbit Alexa647 (1:800; Invitrogen) or Atto 647N (Sigma Aldrich) for STED experiments were incubated in blocking solution for 1 h. Then, cells were washed and mounted with fluoromont-G (Electron Microscopy Sciences). All the procedures were carried out at room temperature. For p-AKt experiments astrocytes were imaged with a Confocal Leica DMI6000 microscope (Leica) and the image analysis was done with ImageJ software (NIH, USA).

### STED imaging and quantification

Astrocytes were processed for immunocytochemistry as described above and imaged using a LEICA SP8 WLL2 on an inverted stand DMI6000 (Leica Microsystems, Mannheim, Germany) and objective HC PL APO 100X oil STED NA 1.40. The depletion lasers used were 660 nm and 775 nm. For overlap analysis, average+5*std from the entire control images was considered as background and the correspondent threshold applied to all images. Then the overlap area over total mitochondria area (Tom20) was calculated. For distance map analysis a threshold equivalent to average+3*std of the control condition was applied to all images and “Distance Map” function from ImageJ software (NIH, USA) used. The resulting images and the data obtained grouped every 0,126 μm (6*pixel size). All image analysis was done with ImageJ software (NIH, USA).

### Electron microscopy

Astrocytes were fixed for 45 min at 4 °C in 4% glutaraldehyde in PBS, washed and fixed again 1 h at room temperature in 2% osmium tetroxide in PBS containing 15 mg/ml of K_4_Fe(CN)_6_. Dehydration was performed with ethanol series (50%, 70%, 95% and absolute ethanol). Thereafter, the samples were embedded progressively in Epon resin. Ultrathin sections were contrasted with 1% lead citrate in water for 1 min. Sample imaging was performed using a Hitachi H7650 transmission microscope operated at 80 KV with a camera Gatan—11 MPx at electron microscopy unit (Bordeaux imaging center). For data analysis mitochondria and ER were manually selected and interfaces between ER and mitochondria were analysed using the Image J plugin [37] (MitoCare). Mitochondria were considered with close ER if their calcium score was bigger than 5.

### Live cell imaging and data analysis

Calcium imaging (Mito-GCaMP6s and RCaMP2) in cortical astrocytes was recorded using an inverted Leica DMI6000 microscope (Leica Microsystems, Wetzlar, Germany) equipped with a confocal Scanner Unit CSU-X1 (Yokogawa Electric Corporation, Tokyo, Japan) and a Evolve EMCCD camera (Photometrics, Tucson, USA). The diode lasers used were at 491 nm and 561 nm and the objective was a HCX PL Apo CS 63× oil 1.4 NA. The 37 °C atmosphere during time-lapse image acquisition was created with an incubator box and an air heating system (Life Imaging Services, Basel, Switzerland). This system was controlled by MetaMorph software (Molecular Devices, Sunnyvale, USA). Only one focal plane was manually selected and recorded every 5 s using an auto-focus system to avoid movement artifacts. Culture media supplemented with 25 μM HEPES was perfused continuously and the different compounds added to the media when desired. On the basis of preliminary experiments WIN was added 15 min after the starting of the recording and ATP 20 min after WIN.

Image processing, analysis and editing were done with ImageJ software (NIH, USA). RCaMP2 measurement linearly adjusted to correct the bleaching of the sample. The signal was averaged every 30 s and the corrected calcium signal (ΔF/F0) was calculated. For both, mitochondrial and cytosolic calcium signals, F0 corresponds to 1 min average before the WIN or ATP application and the area under the curve (AUC) was quantified 4 min after. ATP null responses (except during the irreversible thapsigargin treatment) were considered criteria of exclusion.

### Fluorescence immunohistochemistry

Mice were anaesthetized with chloral hydrate (400 mg*kg−1 body weight) and transcardially perfused with phosphate-buffered solution (PB 0.1 M, pH 7.4) followed by 4% formaldehyde dissolved in PBS (0.1 M, pH 7.4). After perfusion, the brains were removed and incubated several additional hours in the same fixative and finally the brain were embedded with sucrose 30% for 3 days, frozen and kept at −80°C degrees. Serial brain coronal cryosections were cut at 40 μm and collected in 0.1 M PB (pH 7.4) at room temperature (RT).

#### Immunofluorescence NeuN/GFP

Sections were pre-incubated in a blocking solution of 10% donkey serum, 0.02% sodium azide and 0.3% triton X-100 prepared in PBS for 30 min–1 h at room temperature. Free-floating sections were incubated overnight (4 °C) with mouse anti-NeuN (1:500; Millipore, MAB377), rabbit anti-GFP (1:1000; Invitrogen, A11122). The antibodies were prepared in 10% donkey serum in PBS containing 0.02% sodium azide and 0.3% triton X-100. Then, the sections were washed in PBS for 30 min at room temperature. The tissue was incubated with fluorescent anti-rabbit Alexa488, anti-mouse Alexa546 (1:500, Invitrogen) for 2 h and washed in PBS at room temperature. Finally, sections were washed, mounted, dried and a coverslip was added on top with DPX (Fluka Chemie AG). The slides were analysed with Confocal Leica DMI6000 microscope or epifluorescence Leica DM6000 microscope (Leica).

#### Immunofluorescence S100β/GFP

Sections were blocked with 3 % H_2_O_2_ during 30 min. Free-floating sections were incubated overnight (4 °C) with rabbit anti-GFP (1:1000; Invitrogen, A11122), mouse anti-S100β (1:500; Sigma Aldrich, AMAB91038). The antibodies were prepared in 10% donkey serum in PBS containing 0.02% sodium azide and 0.3% triton X-100. Then, the sections were washed in PBS for 30 min at room temperature. The tissue was incubated with fluorescent anti-mouse HRP (1:500, Cell Signaling) and anti-rabbit Alexa488 (1:500, Invitrogen) for 2 h and washed in PBS at room temperature. After, section were incubated with TSA Cy3 (1:300, 10 min, Perkin Elmer). Finally, sections were washed, mounted, dried and a coverslip was added on top with DPX (Fluka Chemie AG). The slides were analysed with Confocal Leica DMI6000 microscope or epifluorescence Leica DM6000 microscope (Leica).

#### Immunofluorescence GFAP

Sections were pre-incubated in a blocking solution of 10% donkey serum, 0.02% sodium azide and 0.3% triton X-100 prepared in PBS for 30 min–1 h at room temperature. Free-floating sections were incubated overnight (4 °C) with rabbit anti-GFAP (1:500; DAKO, Z0334). The antibody was prepared in 10% donkey serum in PBS containing 0.02% sodium azide and 0.3% triton X-100. Then, the sections were washed in PBS for 30 min at room temperature. The tissue was incubated with fluorescent anti-rabbit Alexa 647 (1:500, Invitrogen) for 2 h and washed in PBS at room temperature. Finally, sections were washed, mounted, dried and a coverslip was added on top with DPX (Fluka Chemie AG). The slides were analysed with Confocal Leica DMI6000 microscope or epifluorescence Leica DM6000 microscope (Leica).

### Surgery and AAV administration

Mice (7–10 weeks of age) were anaesthetized by isoflurane or, for two-photon experiments, with an intraperitoneal (i.p.) injection of a mix containing medetomidine (sededorm, 0.27 mg/kg) and midazolam (Dormicum, 5 mg/kg) in sterile NaCl 0.9% (MMF-mix). Analgesia was achieved by local application of 100 μl of lidocaine (lurocaine, 1%) and subcutaneous (s.c.) injection of buprenorphine (buprécare, 0.05 mg/kg). For two-photon experiments, mice additionally administrated with 40 μl of dexamethasone (dexadreson, 0.1 mg/ml) intramuscularly (i.m.) in the quadriceps to reduce the cortical stress during the surgery and to prevent inflammation potentially caused by the friction of the drilling. A heating-pad was positioned underneath the animal to keep the body temperature at 37 °C. Eye dehydration was prevented by topical application of ophthalmic gel. The skin above the skull was shaved with a razor and disinfected with modified ethanol 70% and betadine before an incision was made. Mice were placed into a stereotaxic apparatus (David Kopf Instruments) with mouse adaptor and lateral ear bars.

Cranial window implantation and AAV delivery for two-photon experiments were made as previously described [68, 69]. Briefly, the bregma and lambda were aligned (x and z) after skull’s exposure a ~5 mm custom plastic chamber was attached on the area of interest and a 3 mm craniotomy was made on the right hemisphere above the primary somatosensory cortex, with a pneumatic dental drill (BienAir Medical Technologies, AP-S001), leaving the dura intact. The stereotaxic injections were targeted to the layer 2/3 of the barrel cortex (from bregma: anterior–posterior – 1.5; medial–lateral ± 2.5). 200 nl of virus AAV-GFAP-Mito-GCaMP6s were injected at a maximum rate of 60 nl/min, using a glass pipette (Wiretrol, Drummond) attached to an oil hydraulic manipulator (MO-10, Narishige). After injections, the virus was allowed to diffuse for at least 30 min before the pipette was withdrawn. The craniotomy was covered with sterile saline (0.9% NaCl) and sealed with a 3 mm glass cover slip after viral injection. The chamber, the cover slip and a custom-made stainless steel head stage were well attached to the skull using dental acrylic and dental cement (Jet Repair Acrylic, Lang Dental Manufacturing). Mice were then waked-up by a sub-cutaneous injection of a mixture containing atipamezole (revertor, 2.5 mg/kg), flumazenil (0.5 mg/kg), and buprenorphine (buprécare, 0.1 mg/kg) in sterile NaCl 0.9%. For other viral intrahippocampal and intracortical AAV delivery, AAV vectors were injected with the help of a glass pipette attached to a Nanoject III (Drummond, Broomall, USA). Mice were injected 0.4 μl per injection site at a rate of 0.3 μl per min with the following coordinates: posterior hippocampus, anterior–posterior – 2.6; medial–lateral ± 2.15; dorsal–ventral − 2.0 and − 1.5 (for electrophysiology recordings); anterior hippocampus (anterior–posterior – 2; medial– lateral ± 1; dorsal–ventral − 2.0 (for fiber photometry experiments) and motor cortex: anterior–posterior − 1; medial–lateral ± 1; dorsal–ventral – 0.5 (for fiber photometry). Following virus delivery, the syringe was left in place for 3 min before being slowly withdrawn from the brain. For fiber photometry experiments, the optical fiber (400 μm diameter) was placed 250 μm above the injection site. Mito-GCaMP6s or dnMCU expression was verified by fluorescent immunohistochemistry (see above). Animals were used for experiments 2-4 weeks after injections before experiments.

### 2-photon laser-scanning microscope (2PSLM)-based calcium imaging and analysis

2 weeks after the surgery, anesthetized mice (isoflurane 2%, body temperature maintained at 37°C, and eyes protected from dehydration) were imaged longitudinally using an *in vivo* non-descanned FemtoSmart 2PLSM (Femtonics, Budapest, Hungary) equipped with resonant scanners (operating at 37 Hz) and a ×16 objective (0.8 NA, Nikon). The MES Software (MES v.4.6; Femtonics, Budapest, Hungary) was used to control the microscope, the acquisition parameters, and the TTL-driven synchronization. Mito-GcaMP6s was excited using a Ti:sapphire laser operating at λ=910 nm (Mai Tai DeepSee, Spectra-Physics) with an average excitation power at the focal point lower than 50 mW. Fluorescence from mitochondria expressing GcaMP6s was collected from the same field of view (400 × 400 μm) for 3 consecutive days (one condition per day) for 45 min without injection (day 1), before and after the injection of vehicle (day 2), and before/after the injection of THC (day 3). Manual intraperitoneal injection of either vehicle or THC solution were performed under infrared camera control (Imetronic) during imaging using a custom made tubing system and a needle inserted in the animal prior to recording.

Images from an individual 45 min acquisition session were first laterally realigned so as to compensate for slow, micrometric drift of the field of view. At full temporal resolution (37 Hz), we first used a method for correcting lateral translation with sub-pixel accuracy [70] which works by matching individual images to a dynamically changing template image. We then binned data to 1 Hz to enhance signal-to-noise ratio and re-run the alignment procedure to improve on long-term stability. For each pixel, a time-series of fluorescence fluctuations (*ΔF/F*_*0*_) was then computed. For each time point *t*, *F* was computed as the median fluorescence signal from *t* to *t*+30 s, *F*_*0*_ was the median fluorescence from *t*-30 s to *t*, and *ΔF/F*_*0*_ = *(F* – *F*_0_)/F_0._ Large increase of pixel-wise fluorescence was initially detected based on the criteria *ΔF/F*_*0*_ > 5%. In order to discard spurious increase events that could not be accounted by mitochondrial events (eg noise), we performed morphological opening with a disk of 1-pixel radius of the binary event images (in effect removing isolated pixels too small to be mitochondria). We then grouped clustered positive pixels in the 2D+T stack of images by connected component analysis in 3D (18-connected neighborhood) in order to obtain the final list of temporally and spatially localized mitochondrial fluorescence events. Data sets from different imaging sessions were temporally re-aligned by using the time of i.p. injection as a reference point. For Figure 3F, in each session, we counted the number of fluorescence increase events per 5 min temporal bin and normalized these counts by that of events during the 10 min period immediately preceding injection (Baseline) so as to compensate for injection-independent inter-session and inter-animal variability.

### Fiber photometry imaging and data analysis

3-4 weeks after the surgery freely-moving mice were habituated to handling and the day after imaged using 470 and 405 nm LEDs to excite Mito-GCaMP6s in order to measure mitochondrial calcium activity. The emitted fluorescence is proportional to the calcium concentration for stimulation at 470 nm [71, 72]. The isosbestic stimulation (UV light, 405 nm) was also used in alternation with the blue light (470 nm) to treat the signal after, as the fluorescence emitted after this stimulation is not depending on calcium [73]. The GCaMP6s fluorescence from the astrocytes was collected with a sCMOS camera through an optic fiber divided in 2 sections: a short fiber implanted in the brain of the mouse and a long fiber (modified patchcord), both connected through a ferrule-ferrule (1.25 mm) connection. Matlab program (Matlabworks) was used to synchronize each image recording made by the camera, and the Mito-GCaMP6s light excitation made by the LEDs (470 and 405 nm). The two wavelengths of 470 and 405 nm at a power of 0.1 mW were alternated at a frequency of 20 Hz each (40 Hz alternated light stimulations).

To calculate fluorescence due specifically to fluctuations in calcium and to remove bleaching and movement artifacts, a subtraction of the isosbestic 405 nm signal to the calcium signal (470 nm) was performed. A sliding window average of 2 min was used and ratiometric calcium signal calculated (*ΔF/F*_*0*_). Then calcium transients were detected on the filtered trace (high filter) using a threshold to identify them (2MAD of the baseline before injection, Figure 3J). The first minute previous and after injection were removed from quantification to exclude injection effects and transients were divided in two equal periods of near 15 min each. All data processing was performed using custom MATLAB scripts.

### Hippocampal slice preparation

Animals were decapitated and the brain was rapidly removed and placed in ice-cold cutting solution that contained (in mM): sucrose 180, KCl 2.5, NaH_2_PO_4_ 1.25, NaHCO_3_ 26, MgCl_2_ 12, CaCl_2_ 0.2, glucose 11, and was gassed with 95% O_2_/5% CO_2_ (pH = 7.3–7.4). Slices 350 μm thick were made with a vibratome (VT1200S, Leica) and incubated in ACSF at 34°C. After 30 min slices were kept at room temperature. ACSF contained (in mM): NaCl 123, KCl 2.5, NaH_2_PO_4_ 1.25, NaHCO_3_ 26, MgCl_2_ 1.3, CaCl_2_ 2.5, and glucose 11, and was gassed with 95% O_2_/5% CO_2_ (pH = 7.3–7.4). Slices were transferred to an immersion recording chamber, superfused at 2 ml/min with gassed ACSF and visualized under an Olympus microscope (Olympus Optical).

### Electrophysiology

Electrophysiological simultaneous recordings from 2 CA1 pyramidal neurons (> 60 μm apart) were made in whole-cell configuration of the patch-clamp technique. Patch electrodes had resistances of 3–8 MΩ when filled with the internal solution containing (in mM): KCl 130, HEPES 10, EGTA 1, MgCl2 2, CaCl2 0.3, Phosphocreatine 7, Mg-ATP 3, Na-GTP 0.3 (pH = 7.3, 290 mOsm). Recordings were obtained with a MultiClamp 700B amplifier (Molecular devices). Membrane potential was held at −70 mV and series and input resistances were monitored throughout the experiment using a −5 mV pulse. Cells were discarded when series and input resistances changed >20%. Signals were fed to a PC through a DigiData 1440A interface board. Signals were filtered at 1 KHz and acquired at 10 KHz sampling rate. The pCLAMP 10.7 (Axon instruments) software was used for stimulus generation, data display, acquisition and storage. Excitatory postsynaptic currents (EPSCs) were isolated in the presence of picrotoxin (50 μM) and CGP54626 (1 μM) to block GABAA and GABAB receptors, respectively. All recordings were performed at 34°C.

### Synaptic stimulation

Theta capillaries filled with ACSF were placed in the Stratum Radiatum for bipolar stimulation of Schaffer collaterals. Paired pulses (2 ms duration with 50 ms interval) were continuously delivered at 0.33 Hz using a Digitimer Ltd stimulator. Stimulus intensity (0.1–10 mA) was adjusted to meet the conditions that putatively stimulate single or very few presynaptic fibers [74–76]. Synaptic parameters analysed were: success rate (ratio between the number of successes versus total number of stimuli) and EPSC amplitude (mean peak amplitude of the successes, excluding failures). A response was considered a failure if the amplitude of the current was <3 times the standard deviation of the baseline current and was verified by visual inspection.

During the neuronal depolarization (ND) protocol one pyramidal neuron was depolarized to 0 mV for 5 s to stimulate endocannabinoid release [15, 31, 77–80]. To illustrate the time course of the effects of ND, synaptic parameters were grouped in 60 s bins. To determine synaptic changes, mean EPSCs (n = 60 EPSCs) recorded 3 min before the stimulus (basal) were compared with mean EPSCs (n = 20 EPSCs) recorded 1 min after ND. The presence of synaptic potentiation was determined to occur when the success rate increased more than two standard deviations from the baseline.

### Confocal acquisition of ND-induced astroglial calcium events and data analysis

Neuronal depolarization (to 0 mV, 5s) was made using a Patchmaster software and an EPC10 amplifier (HEKA Elektronik, Lambrecht/Pfalz, Germany). Confocal acquisition were performed using a Leica SP5 on an upright stand DM6000 (Leica Microsystems, Mannheim, Germany) and with a 20× (NA 1) water immersion objective. Images were acquired every 0.55-0.66 s using a 488 nm Argon laser and a 543 nm Helium/Neon laser for stimulation of GCaMP6f and SR101 intensities, respectively. Pixel size was 0.55 μm.

Image analysis was done using a custom Image J macro (https://github.com/Romanserrat/3D-calcium-counting). Briefly, images were first laterally realigned so as to compensate for slow, micrometric drift of the field of view. Then, calcium events (above mean+4*std) were isolated and selected in 3 dimensions (x, y, t) using 3D object counter and 3D viewer Image J plugins (events bigger than 40 pixels). From each calcium event, the amplitude, frequency, duration and spreading coefficient (max x distance * max y distance) were calculated. For quantification purposes only the mean values (per slice) of the calcium events starting 30 s before or 30 s after neuronal depolarization were considered.

### Statistical analyses

All graphs and statistical analyses were performed using GraphPad software (version 5.0 or 6.0). Results were expressed as means of independent data points ± s.e.m. Parametric test was used for the data and analysed using the appropriate statistical test (detailed statistical data for each experiment are reported in Table S1,2).

## References

1. Allaman, I., Belanger, M., and Magistretti, P.J. (2011). Astrocyte-neuron metabolic relationships: for better and for worse. Trends in neurosciences 34, 76–87.

2. Haydon, P.G., and Carmignoto, G. (2006). Astrocyte control of synaptic transmission and neurovascular coupling. Physiological reviews 86, 1009–1031.

3. Perea, G., Navarrete, M., and Araque, A. (2009). Tripartite synapses: astrocytes process and control synaptic information. Trends in neurosciences 32, 421–431.

4. Perez-Alvarez, A., and Araque, A. (2013). Astrocyte-neuron interaction at tripartite synapses. Current drug targets 14, 1220–1224.

5. Perea, G., and Araque, A. (2005). Synaptic regulation of the astrocyte calcium signal. Journal of neural transmission 112, 127–135.

6. Scemes, E., and Giaume, C. (2006). Astrocyte calcium waves: what they are and what they do. Glia 54, 716–725.

7. Volterra, A., Liaudet, N., and Savtchouk, I. (2014). Astrocyte Ca(2)(+) signalling: an unexpected complexity. Nature reviews. Neuroscience 15, 327–335.

8. Agarwal, A., Wu, P.H., Hughes, E.G., Fukaya, M., Tischfield, M.A., Langseth, A.J., Wirtz, D., and Bergles, D.E. (2017). Transient Opening of the Mitochondrial Permeability Transition Pore Induces Microdomain Calcium Transients in Astrocyte Processes. Neuron 93, 587–605 e587.

9. Rizzuto, R., De Stefani, D., Raffaello, A., and Mammucari, C. (2012). Mitochondria as sensors and regulators of calcium signalling. Nature reviews. Molecular cell biology 13, 566–578.

10. Simpson, P.B., Mehotra, S., Lange, G.D., and Russell, J.T. (1997). High density distribution of endoplasmic reticulum proteins and mitochondria at specialized Ca2+ release sites in oligodendrocyte processes. The Journal of biological chemistry 272, 22654–22661.

11. Stephen, T.L., Higgs, N.F., Sheehan, D.F., Al Awabdh, S., Lopez-Domenech, G., Arancibia-Carcamo, I.L., and Kittler, J.T. (2015). Miro1 Regulates Activity-Driven Positioning of Mitochondria within Astrocytic Processes Apposed to Synapses to Regulate Intracellular Calcium Signaling. The Journal of neuroscience : the official journal of the Society for Neuroscience 35, 15996–16011.

12. Rizzuto, R., Brini, M., Murgia, M., and Pozzan, T. (1993). Microdomains with high Ca2+ close to IP3-sensitive channels that are sensed by neighboring mitochondria. Science 262, 744–747.

13. Gӧbel, J., Engelhardt, E., Pelzer, P., Sakthivelu, V., Jahn, H.M., Jevtic, M., Folz-Donahue, K., Kukat, C., Schauss, A., Frese, C.K., et al. (2020). Mitochondria-Endoplasmic Reticulum Contacts in Reactive Astrocytes Promote Vascular Remodeling. Cell Metab 31, 791–808.e798.

14. Gomez-Gonzalo, M., Navarrete, M., Perea, G., Covelo, A., Martin-Fernandez, M., Shigemoto, R., Lujan, R., and Araque, A. (2015). Endocannabinoids Induce Lateral Long-Term Potentiation of Transmitter Release by Stimulation of Gliotransmission. Cerebral cortex 25, 3699–3712.

15. Navarrete, M., and Araque, A. (2010). Endocannabinoids potentiate synaptic transmission through stimulation of astrocytes. Neuron 68, 113–126.

16. Covelo, A., and Araque, A. (2016). Lateral regulation of synaptic transmission by astrocytes. Neuroscience 323, 62–66.

17. Covelo, A., and Araque, A. (2018). Neuronal activity determines distinct gliotransmitter release from a single astrocyte. eLife 7.

18. Busquets-Garcia, A., Bains, J., and Marsicano, G. (2018). CB1 Receptor Signaling in the Brain: Extracting Specificity from Ubiquity. Neuropsychopharmacology : official publication of the American College of Neuropsychopharmacology 43, 4–20.

19. Piomelli, D. (2003). The molecular logic of endocannabinoid signalling. Nature reviews. Neuroscience 4, 873–884.

20. Zou, S., and Kumar, U. (2018). Cannabinoid Receptors and the Endocannabinoid System: Signaling and Function in the Central Nervous System. International journal of molecular sciences 19.

21. Benard, G., Massa, F., Puente, N., Lourenco, J., Bellocchio, L., Soria-Gomez, E., Matias, I., Delamarre, A., Metna-Laurent, M., Cannich, A., et al. (2012). Mitochondrial CB(1) receptors regulate neuronal energy metabolism. Nature neuroscience 15, 558–564.

22. Hebert-Chatelain, E., Desprez, T., Serrat, R., Bellocchio, L., Soria-Gomez, E., Busquets-Garcia, A., Pagano Zottola, A.C., Delamarre, A., Cannich, A., Vincent, P., et al. (2016). A cannabinoid link between mitochondria and memory. Nature 539, 555–559.

23. Jimenez-Blasco, D., Busquets-Garcia, A., Hebert-Chatelain, E., Serrat, R., Vicente-Gutierrez, C., Ioannidou, C., Gomez-Sotres, P., Lopez-Fabuel, I., Resch-Beusher, M., Resel, E., et al. (2020). Glucose metabolism links astroglial mitochondria to cannabinoid effects. Nature.

24. Koch, M., Varela, L., Kim, J.G., Kim, J.D., Hernandez-Nuno, F., Simonds, S.E., Castorena, C.M., Vianna, C.R., Elmquist, J.K., Morozov, Y.M., et al. (2015). Hypothalamic POMC neurons promote cannabinoid-induced feeding. Nature 519, 45–50.

25. Ma, L., Jia, J., Niu, W., Jiang, T., Zhai, Q., Yang, L., Bai, F., Wang, Q., and Xiong, L. (2015). Mitochondrial CB1 receptor is involved in ACEA-induced protective effects on neurons and mitochondrial functions. Scientific reports 5, 12440.

26. Xu, Z., Lv, X.A., Dai, Q., Ge, Y.Q., and Xu, J. (2016). Acute upregulation of neuronal mitochondrial type-1 cannabinoid receptor and it’s role in metabolic defects and neuronal apoptosis after TBI. Molecular brain 9, 75.

27. Gutierrez-Rodriguez, A., Bonilla-Del Rio, I., Puente, N., Gomez-Urquijo, S.M., Fontaine, C.J., Egana-Huguet, J., Elezgarai, I., Ruehle, S., Lutz, B., Robin, L.M., et al. (2018). Localization of the cannabinoid type-1 receptor in subcellular astrocyte compartments of mutant mouse hippocampus. Glia 66, 1417–1431.

28. Han, J., Kesner, P., Metna-Laurent, M., Duan, T., Xu, L., Georges, F., Koehl, M., Abrous, D.N., Mendizabal-Zubiaga, J., Grandes, P., et al. (2012). Acute cannabinoids impair working memory through astroglial CB1 receptor modulation of hippocampal LTD. Cell 148, 1039–1050.

29. Oliveira da Cruz, J.F., Robin, L.M., Drago, F., Marsicano, G., and Metna-Laurent, M. (2016). Astroglial type-1 cannabinoid receptor (CB1): A new player in the tripartite synapse. Neuroscience 323, 35–42.

30. Hegyi, Z., Olah, T., Koszeghy, A., Piscitelli, F., Hollo, K., Pal, B., Csernoch, L., Di Marzo, V., and Antal, M. (2018). CB1 receptor activation induces intracellular Ca(2+) mobilization and 2-arachidonoylglycerol release in rodent spinal cord astrocytes. Scientific reports 8, 10562.

31. Navarrete, M., and Araque, A. (2008). Endocannabinoids mediate neuron-astrocyte communication. Neuron 57, 883–893.

32. Inoue, M., Takeuchi, A., Horigane, S., Ohkura, M., Gengyo-Ando, K., Fujii, H., Kamijo, S., Takemoto-Kimura, S., Kano, M., Nakai, J., et al. (2015). Rational design of a high-affinity, fast, red calcium indicator R-CaMP2. Nature methods 12, 64–70.

33. Bowser, D.N., and Khakh, B.S. (2007). Vesicular ATP is the predominant cause of intercellular calcium waves in astrocytes. The Journal of general physiology 129, 485–491.

34. Sakuragi, S., Niwa, F., Oda, Y., Mikoshiba, K., and Bannai, H. (2017). Astroglial Ca(2+) signaling is generated by the coordination of IP3R and store-operated Ca(2+) channels. Biochemical and biophysical research communications 486, 879–885.

35. Sherwood, M.W., Arizono, M., Hisatsune, C., Bannai, H., Ebisui, E., Sherwood, J.L., Panatier, A., Oliet, S.H., and Mikoshiba, K. (2017). Astrocytic IP3 Rs: Contribution to Ca(2+) signalling and hippocampal LTP. Glia 65, 502–513.

36. Csordas, G., Weaver, D., and Hajnoczky, G. (2018). Endoplasmic Reticulum-Mitochondrial Contactology: Structure and Signaling Functions. Trends in cell biology 28, 523–540.

37. Bartok, A., Weaver, D., Golenar, T., Nichtova, Z., Katona, M., Bansaghi, S., Alzayady, K.J., Thomas, V.K., Ando, H., Mikoshiba, K., et al. (2019). IP3 receptor isoforms differently regulate ER-mitochondrial contacts and local calcium transfer. Nature communications 10, 3726.

38. Di Marco, G., Vallese, F., Jourde, B., Bergsdorf, C., Sturlese, M., De Mario, A., Techer-Etienne, V., Haasen, D., Oberhauser, B., Schleeger, S., et al. (2020). A High-Throughput Screening Identifies MICU1 Targeting Compounds. Cell reports 30, 2321–2331 e2326.

39. Marchi, S., Corricelli, M., Branchini, A., Vitto, V.A.M., Missiroli, S., Morciano, G., Perrone, M., Ferrarese, M., Giorgi, C., Pinotti, M., et al. (2019). Akt-mediated phosphorylation of MICU1 regulates mitochondrial Ca(2+) levels and tumor growth. The EMBO journal 38.

40. Raffaello, A., De Stefani, D., Sabbadin, D., Teardo, E., Merli, G., Picard, A., Checchetto, V., Moro, S., Szabo, I., and Rizzuto, R. (2013). The mitochondrial calcium uniporter is a multimer that can include a dominant-negative pore-forming subunit. The EMBO journal 32, 2362–2376.

41. Cahoy, J.D., Emery, B., Kaushal, A., Foo, L.C., Zamanian, J.L., Christopherson, K.S., Xing, Y., Lubischer, J.L., Krieg, P.A., Krupenko, S.A., et al. (2008). A transcriptome database for astrocytes, neurons, and oligodendrocytes: a new resource for understanding brain development and function. The Journal of neuroscience : the official journal of the Society for Neuroscience 28, 264–278.

42. Hammond, S.L., Leek, A.N., Richman, E.H., and Tjalkens, R.B. (2017). Cellular selectivity of AAV serotypes for gene delivery in neurons and astrocytes by neonatal intracerebroventricular injection. PloS one 12, e0188830.

43. Pagano Zottola, A.C., Soria-Gomez, E., Bonilla-del-Río, I., Muguruza, C., Terral, G., Robin, L.M., da Cruz, J.F.O., Redon, B., Lesté-Lasserre, T., Tolentino-Cortes, T., et al. (2020). A new mutant mouse model lacking mitochondrial-associated CB1 receptor. bioRxiv, 2020.2003.2030.009472.

44. Markus, N.M., Hasel, P., Qiu, J., Bell, K.F., Heron, S., Kind, P.C., Dando, O., Simpson, T.I., and Hardingham, G.E. (2016). Expression of mRNA Encoding Mcu and Other Mitochondrial Calcium Regulatory Genes Depends on Cell Type, Neuronal Subtype, and Ca2+ Signaling. PloS one 11, e0148164.

45. Celsi, F., Pizzo, P., Brini, M., Leo, S., Fotino, C., Pinton, P., and Rizzuto, R. (2009). Mitochondria, calcium and cell death: a deadly triad in neurodegeneration. Biochimica et biophysica acta 1787, 335–344.

46. Duchen, M.R. (2000). Mitochondria and Ca(2+)in cell physiology and pathophysiology. Cell calcium 28, 339–348.

47. O-Uchi, J., Pan, S., and Sheu, S.S. (2012). Perspectives on: SGP symposium on mitochondrial physiology and medicine: molecular identities of mitochondrial Ca2+ influx mechanism: updated passwords for accessing mitochondrial Ca2+-linked health and disease. The Journal of general physiology 139, 435–443.

48. Santo-Domingo, J., and Demaurex, N. (2010). Calcium uptake mechanisms of mitochondria. Biochimica et biophysica acta 1797, 907–912.

49. Carafoli, E. (2003). Historical review: mitochondria and calcium: ups and downs of an unusual relationship. Trends in biochemical sciences 28, 175–181.

50. Jhun, B.S., Mishra, J., Monaco, S., Fu, D., Jiang, W., Sheu, S.S., and J, O.U. (2016). The mitochondrial Ca2+ uniporter: regulation by auxiliary subunits and signal transduction pathways. American journal of physiology. Cell physiology 311, C67–80.

51. Tarasova, N.V., Vishnyakova, P.A., Logashina, Y.A., and Elchaninov, A.V. (2019). Mitochondrial Calcium Uniporter Structure and Function in Different Types of Muscle Tissues in Health and Disease. International journal of molecular sciences 20.

52. Abadir, P.M., Foster, D.B., Crow, M., Cooke, C.A., Rucker, J.J., Jain, A., Smith, B.J., Burks, T.N., Cohn, R.D., Fedarko, N.S., et al. (2011). Identification and characterization of a functional mitochondrial angiotensin system. Proceedings of the National Academy of Sciences of the United States of America 108, 14849–14854.

53. Belous, A.E., Jones, C.M., Wakata, A., Knox, C.D., Nicoud, I.B., Pierce, J., and Chari, R.S. (2006). Mitochondrial calcium transport is regulated by P2Y1- and P2Y2-like mitochondrial receptors. Journal of cellular biochemistry 99, 1165–1174.

54. Lahuna, O., and Jockers, R. (2018). [Mitochondrial signaling of G protein-coupled receptors]. Biologie aujourd’hui 212, 21–26.

55. Suofu, Y., Li, W., Jean-Alphonse, F.G., Jia, J., Khattar, N.K., Li, J., Baranov, S.V., Leronni, D., Mihalik, A.C., He, Y., et al. (2017). Dual role of mitochondria in producing melatonin and driving GPCR signaling to block cytochrome c release. Proceedings of the National Academy of Sciences of the United States of America 114, E7997–E8006.

56. Williams, G.S., Boyman, L., Chikando, A.C., Khairallah, R.J., and Lederer, W.J. (2013). Mitochondrial calcium uptake. Proceedings of the National Academy of Sciences of the United States of America 110, 10479–10486.

57. Boyman, L., Chikando, A.C., Williams, G.S., Khairallah, R.J., Kettlewell, S., Ward, C.W., Smith, G.L., Kao, J.P., and Lederer, W.J. (2014). Calcium movement in cardiac mitochondria. Biophysical journal 107, 1289–1301.

58. Boitier, E., Rea, R., and Duchen, M.R. (1999). Mitochondria exert a negative feedback on the propagation of intracellular Ca2+ waves in rat cortical astrocytes. The Journal of cell biology 145, 795–808.

59. Jackson, J.G., and Robinson, M.B. (2015). Reciprocal Regulation of Mitochondrial Dynamics and Calcium Signaling in Astrocyte Processes. The Journal of neuroscience : the official journal of the Society for Neuroscience 35, 15199–15213.

60. Reyes, R.C., and Parpura, V. (2008). Mitochondria modulate Ca2+-dependent glutamate release from rat cortical astrocytes. The Journal of neuroscience : the official journal of the Society for Neuroscience 28, 9682–9691.

61. Foskett, J.K., and Daniel Mak, D.O. (2010). Regulation of IP(3)R Channel Gating by Ca(2+) and Ca(2+) Binding Proteins. Current topics in membranes 66, 235–272.

62. Araque, A., Carmignoto, G., and Haydon, P.G. (2001). Dynamic signaling between astrocytes and neurons. Annual review of physiology 63, 795–813.

63. Nedergaard, M., Ransom, B., and Goldman, S.A. (2003). New roles for astrocytes: redefining the functional architecture of the brain. Trends in neurosciences 26, 523–530.

64. Volterra, A., and Meldolesi, J. (2005). Astrocytes, from brain glue to communication elements: the revolution continues. Nature reviews. Neuroscience 6, 626–640.

65. Rizzuto, R., Brini, M., Pizzo, P., Murgia, M., and Pozzan, T. (1995). Chimeric green fluorescent protein as a tool for visualizing subcellular organelles in living cells. Current biology : CB 5, 635–642.

66. Rizzuto, R., Nakase, H., Darras, B., Francke, U., Fabrizi, G.M., Mengel, T., Walsh, F., Kadenbach, B., DiMauro, S., and Schon, E.A. (1989). A gene specifying subunit VIII of human cytochrome c oxidase is localized to chromosome 11 and is expressed in both muscle and non-muscle tissues. The Journal of biological chemistry 264, 10595–10600.

67. McClure, C., Cole, K.L., Wulff, P., Klugmann, M., and Murray, A.J. (2011). Production and titering of recombinant adeno-associated viral vectors. Journal of visualized experiments : JoVE, e3348.

68. Gambino, F., Pages, S., Kehayas, V., Baptista, D., Tatti, R., Carleton, A., and Holtmaat, A. (2014). Sensory-evoked LTP driven by dendritic plateau potentials in vivo. Nature 515, 116–119.

69. Holtmaat, A., Bonhoeffer, T., Chow, D.K., Chuckowree, J., De Paola, V., Hofer, S.B., Hubener, M., Keck, T., Knott, G., Lee, W.C., et al. (2009). Long-term, high-resolution imaging in the mouse neocortex through a chronic cranial window. Nature protocols 4, 1128–1144.

70. Pnevmatikakis, E.A., and Giovannucci, A. (2017). NoRMCorre: An online algorithm for piecewise rigid motion correction of calcium imaging data. Journal of neuroscience methods 291, 83–94.

71. Akerboom, J., Carreras Calderon, N., Tian, L., Wabnig, S., Prigge, M., Tolo, J., Gordus, A., Orger, M.B., Severi, K.E., Macklin, J.J., et al. (2013). Genetically encoded calcium indicators for multi-color neural activity imaging and combination with optogenetics. Frontiers in molecular neuroscience 6, 2.

72. Ohkura, M., Sasaki, T., Sadakari, J., Gengyo-Ando, K., Kagawa-Nagamura, Y., Kobayashi, C., Ikegaya, Y., and Nakai, J. (2012). Genetically encoded green fluorescent Ca2+ indicators with improved detectability for neuronal Ca2+ signals. PloS one 7, e51286.

73. Lutcke, H., Murayama, M., Hahn, T., Margolis, D.J., Astori, S., Zum Alten Borgloh, S.M., Gobel, W., Yang, Y., Tang, W., Kugler, S., et al. (2010). Optical recording of neuronal activity with a genetically-encoded calcium indicator in anesthetized and freely moving mice. Frontiers in neural circuits 4, 9.

74. Allen, C., and Stevens, C.F. (1994). An evaluation of causes for unreliability of synaptic transmission. Proceedings of the National Academy of Sciences of the United States of America 91, 10380–10383.

75. Isaac, J.T., Hjelmstad, G.O., Nicoll, R.A., and Malenka, R.C. (1996). Long-term potentiation at single fiber inputs to hippocampal CA1 pyramidal cells. Proceedings of the National Academy of Sciences of the United States of America 93, 8710–8715.

76. Raastad, M. (1995). Extracellular activation of unitary excitatory synapses between hippocampal CA3 and CA1 pyramidal cells. The European journal of neuroscience 7, 1882–1888.

77. Chevaleyre, V., and Castillo, P.E. (2004). Endocannabinoid-mediated metaplasticity in the hippocampus. Neuron 43, 871–881.

78. Kreitzer, A.C., and Regehr, W.G. (2001). Retrograde inhibition of presynaptic calcium influx by endogenous cannabinoids at excitatory synapses onto Purkinje cells. Neuron 29, 717–727.

79. Ohno-Shosaku, T., Maejima, T., and Kano, M. (2001). Endogenous cannabinoids mediate retrograde signals from depolarized postsynaptic neurons to presynaptic terminals. Neuron 29, 729–738.

80. Wilson, R.I., and Nicoll, R.A. (2001). Endogenous cannabinoids mediate retrograde signalling at hippocampal synapses. Nature 410, 588–592.

