## Supplementary Information for "Astroglial calcium transfer from endoplasmic reticulum to mitochondria determines synaptic integration"

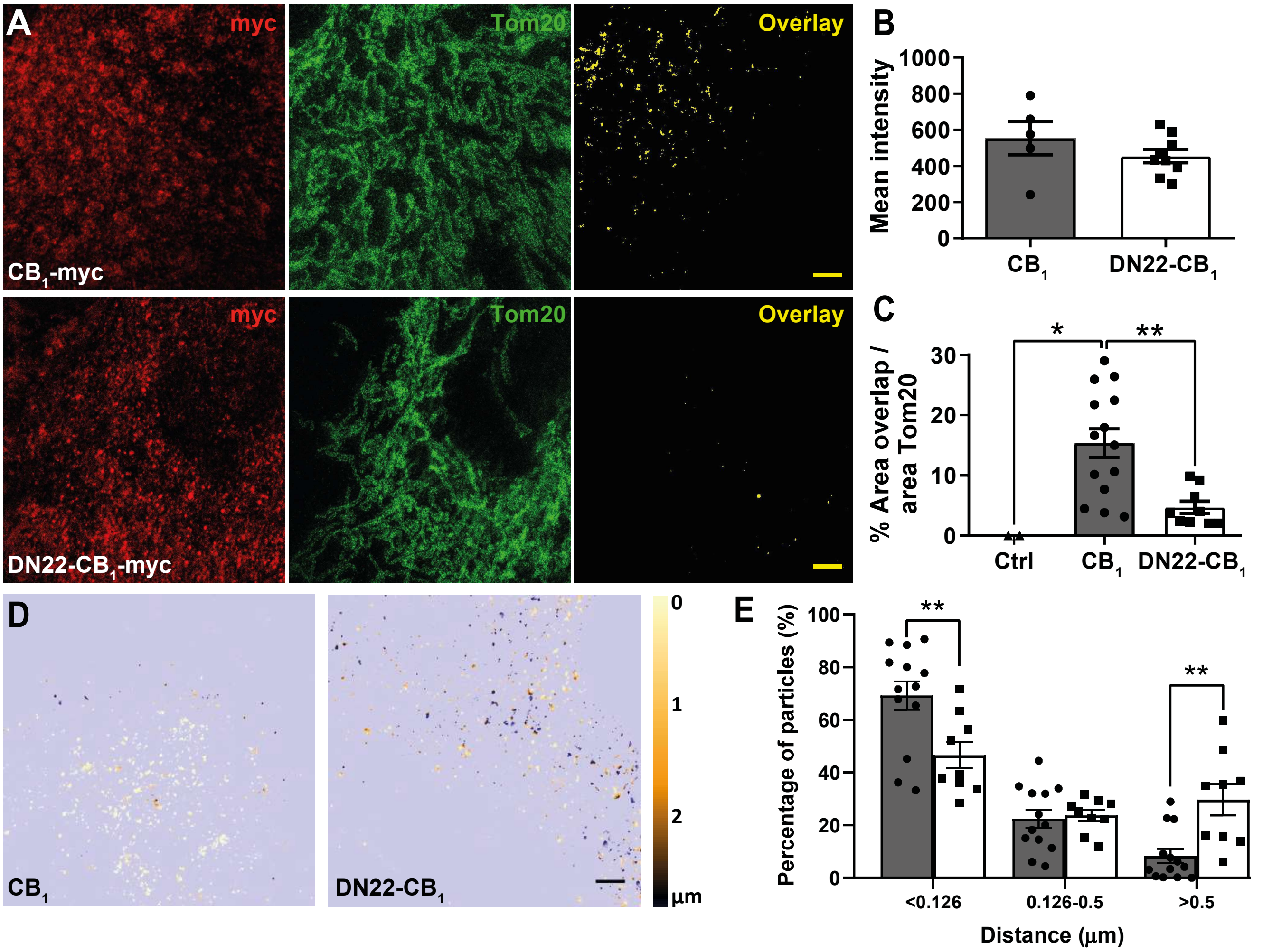


**Figure S1. A** Representative STED images of $CB_1$-myc (upper panels) and DN22-$CB_1$-myc (lower panels) transfected astrocytes. In green, Tom20 (mitochondrial marker) staining; in red, myc ($CB_1$) staining; in yellow, overlay images. Scale bar: 2 µm. **B** Mean intensity quantification of $CB_1$ levels. **C** Percentage of overlap area over Tom20 (total mitochondria). Ctrl: Control, empty-vector transfected cells. **D** Representative distance map images obtained from the STED images used in (C). On the right color-distance legend. Scale bar: 2 µm. **E** Percentage of $CB_1$ particles at given distances. *p-value<0.05, **p-value<0.01.

### Figure S1

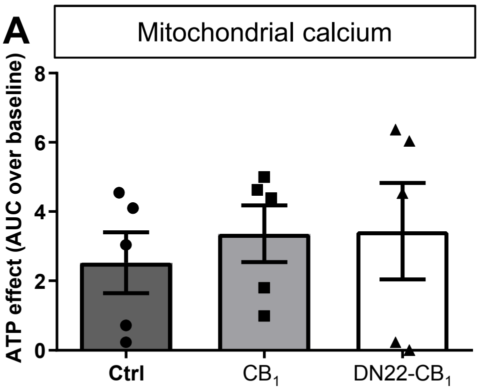


**Figure S2. A** ATP effect on mitochondrial calcium (Area under the curve, AUC) on control (Ctrl), $CB_1$ and DN22-$CB_1$ expressing astrocytes. Baseline: 1 min before ATP-treatment.

### Figure S2

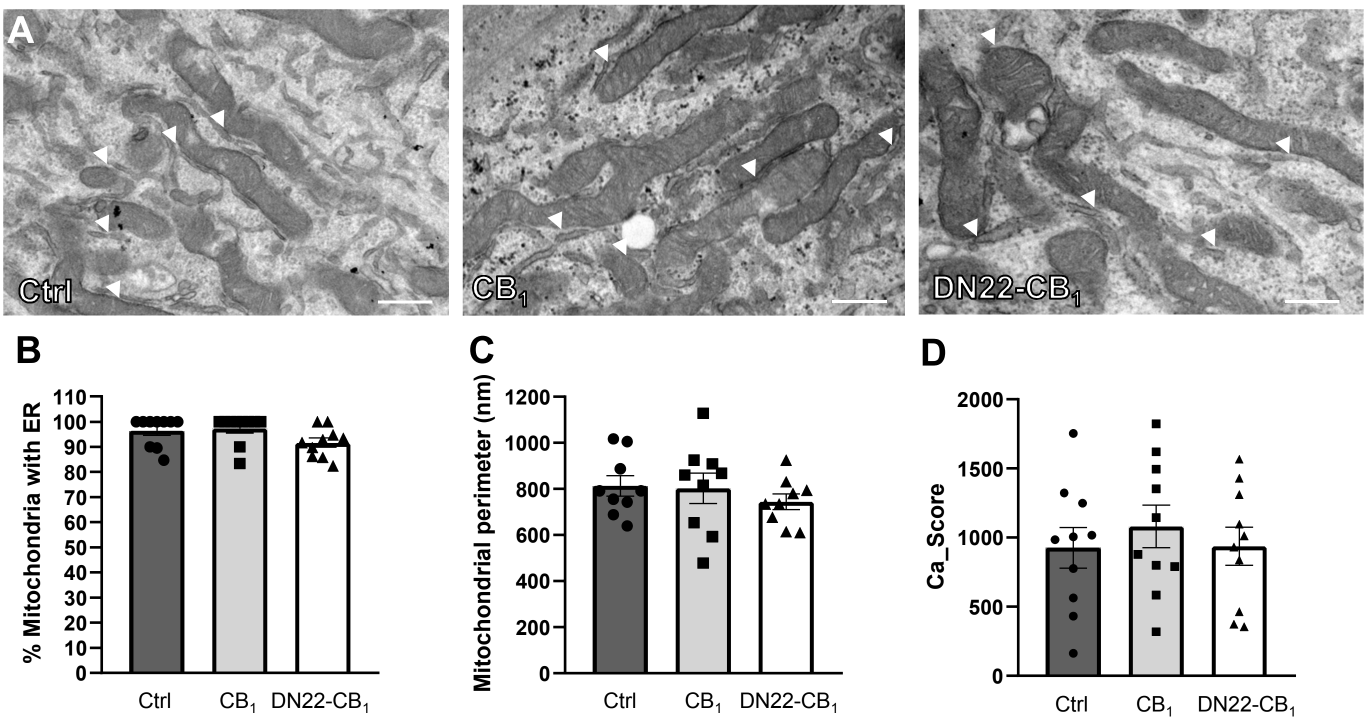


**Figure S3. A** Representative electron microscopy images from control (Ctrl), $CB_1$ and DN22-$CB_1$ transfected astrocytes. White arrow heads: MERCS. Scale bar: 0,5 µm. **B** Percentage of mitochondria with close ER. **C** Mean mitochondrial perimeter per cell. **D** Cell calcium score.

**Figure S3**

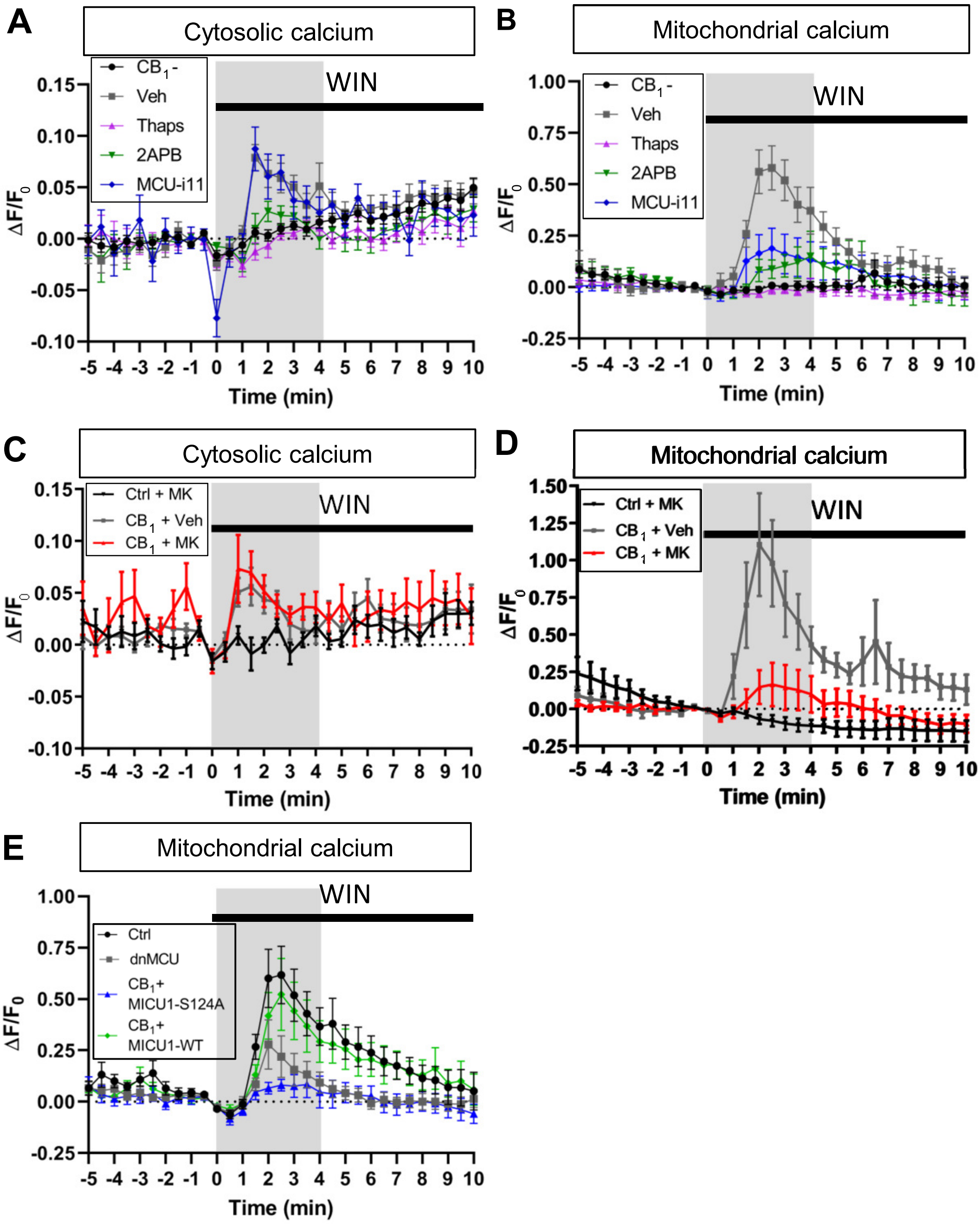


**Figure S4. A-E** $\Delta F/F_0$ traces from RCaMP2 (A,C) or Mito-GCaMP6s (B,D,E) before and after WIN treatment in the different analyzed conditions shown in Figure 2.

### Figure S4

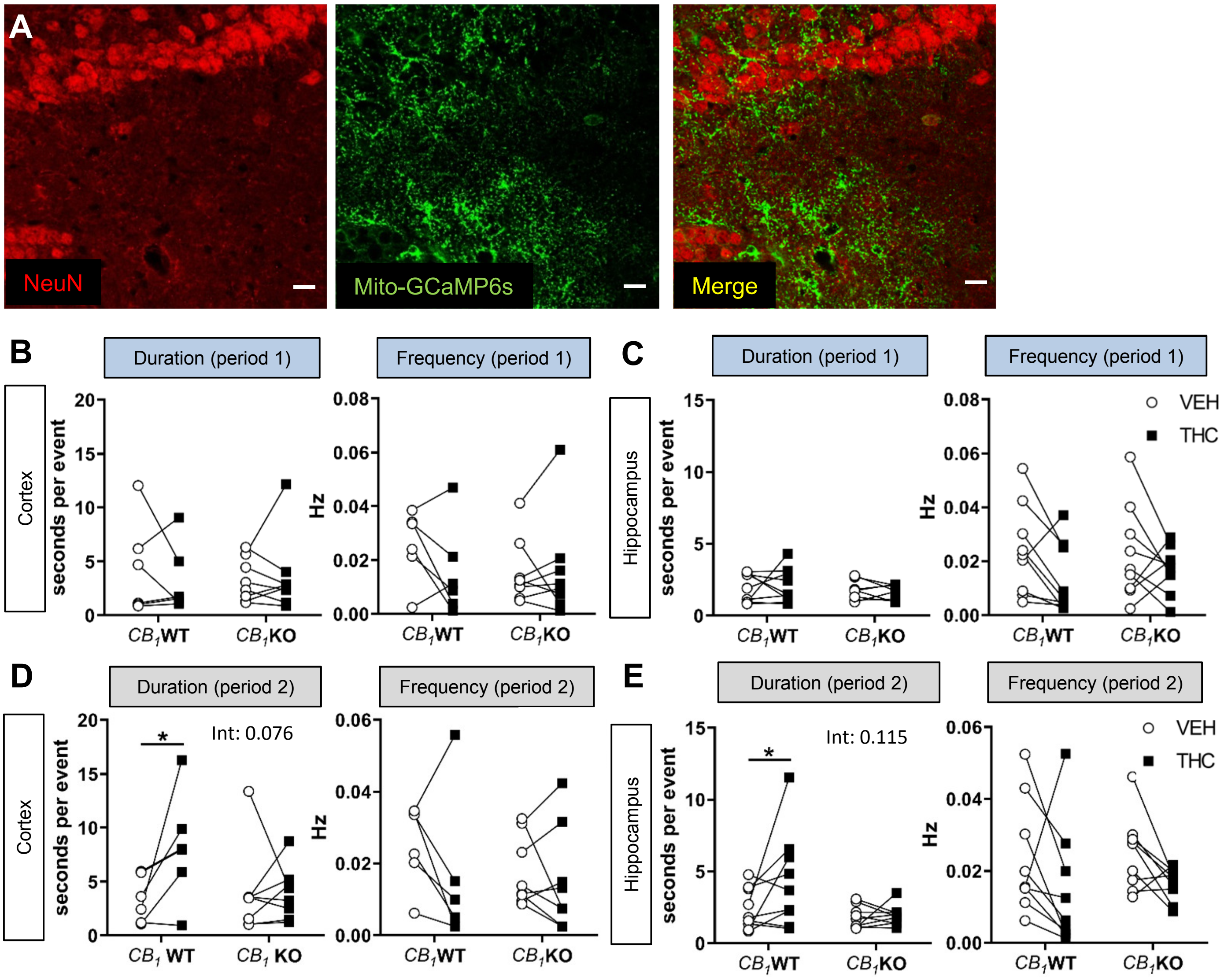


**Figure S5. A** Immunohistochemistry representative images from mouse hippocampi injected with AAV-GFAP-Mito-GCaMP6s (in green). Neuronal marker NeuN in red. Scale bar: 20 µm. **B-E** Cortical (B-D) or hippocampal (C,E) event duration and frequency in *$CB_1$* WT and *$CB_1$* KO mice treated with Vehicle (VEH) or THC during the 2 time periods highlighted in Figure 3J. Int: Paired two-way ANOVA interaction. * p-value<0.05.

### Figure S5

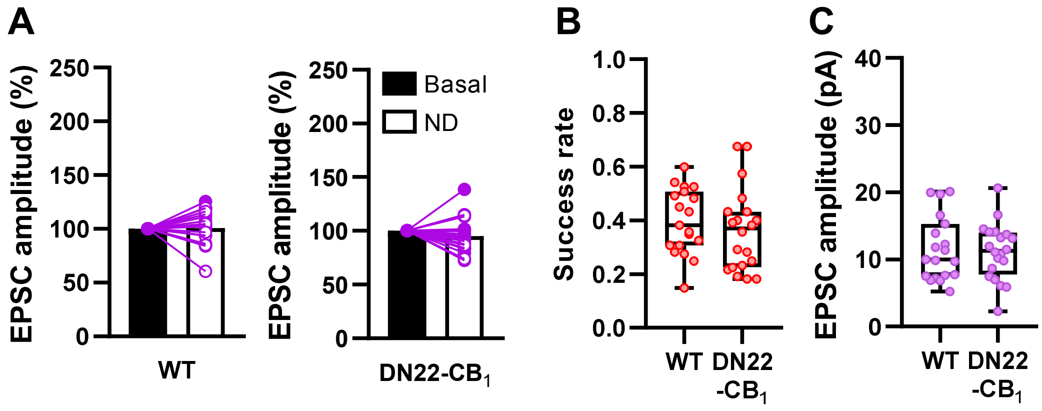


**Figure S6. A** Normalized EPSC amplitude before (Basal) and after neuronal depolarization (ND). White and purple circles represent potentiated and non-potentiated synapses, respectively. **B,C** Basal success rate (B) and EPSC amplitude (C) recorded from WT and DN22-$CB_1$ mice.

### Figure S6

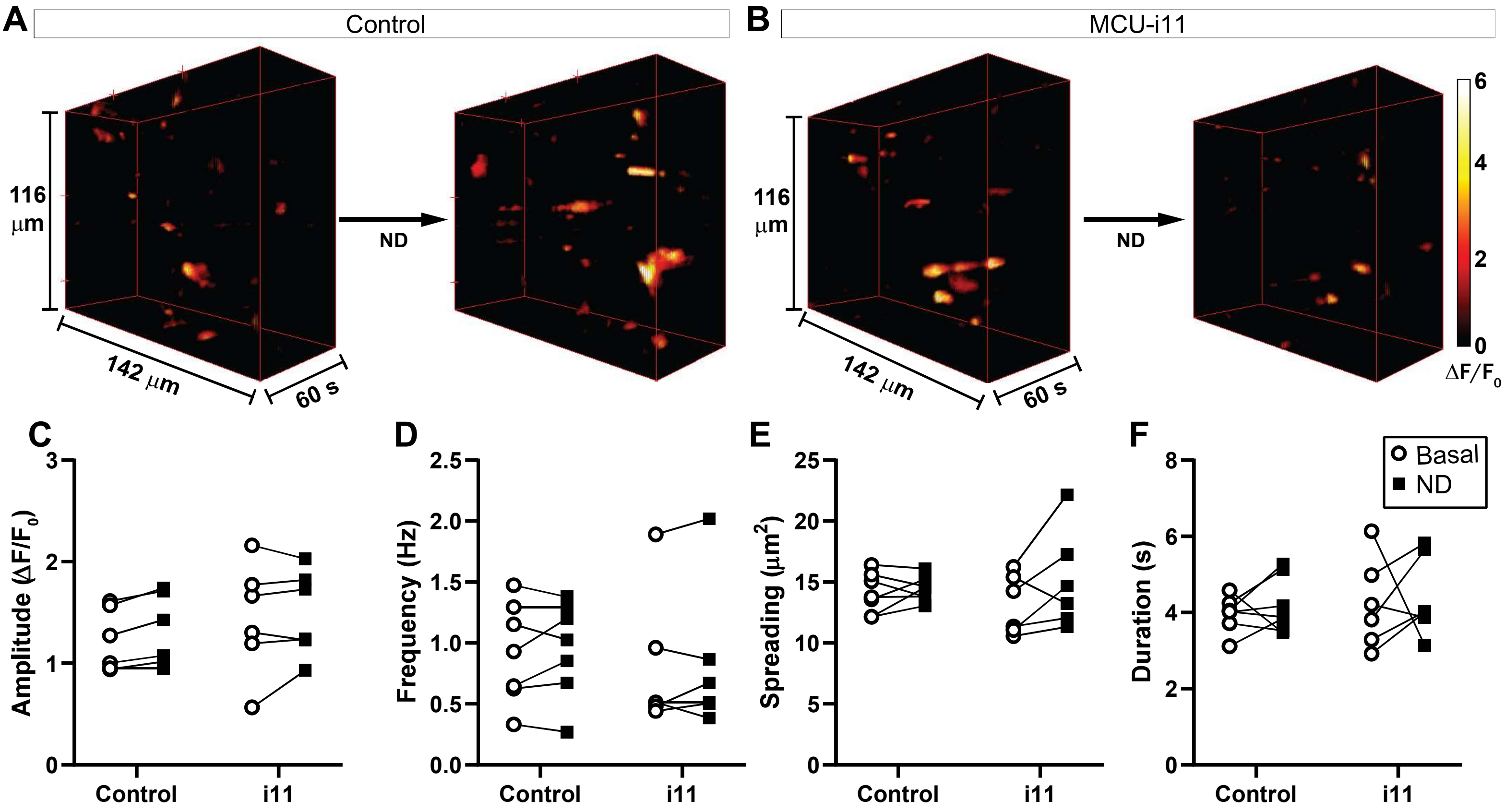


**Figure S7. A,B** 3D representation (x, y and time) of the calcium events occuring before or after neuronal depolarization (ND) in Control condition (A) or after 20min MCU-i11 treatment (B). **C-F** Mean amplitude (C), Frequency (D), mean spreading area (E) and mean duration (F) of the events smaller than 40 μm². ND: neuronal depolarization.

**Figure S7**

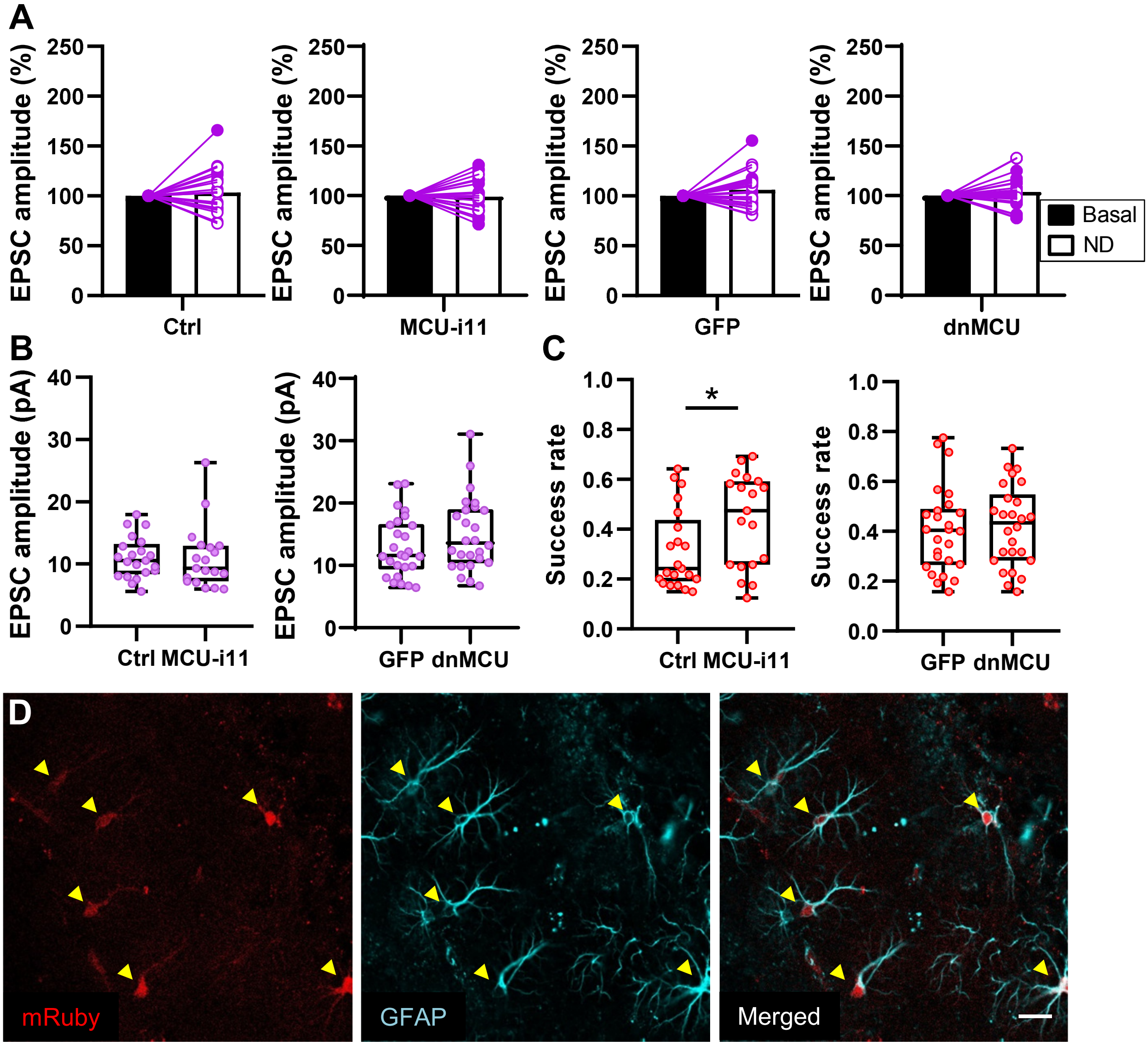


**Figure S8. A** Normalized EPSC amplitude before (Basal) and after neuronal depolarization (ND). White and purple circles represent potentiated and non-potentiated synapses, respectively. **B** EPSC amplitude recorded in the different experimental conditions and transgenic mice. **C** Like B but for basal success rate. **D** Representative **i**mmunohistochemistry from mouse hippocampi slices infected with AAV-GFAP-dnMCU-IRES-mRuby (in red). Astroglial marker GFAP in cyan. Scale bar: 20 μm.

### Figure S8

| Figure | Conditions | "n" | Analysis (post-hoc test) | Factor analyzed | | P value |
|---|---|---|---|---|---|---|
| Figure 1C | WIN on Genotype | 5 | One-way ANOVA (Tukey) | Ctrl vs CB1 vs DN22-CB1 | F (2, 12) = 5.932 | 0.0162 |
| Figure 1F | WIN on Genotype | 5 | One-way ANOVA (Tukey) | Ctrl vs CB1 vs DN22-CB1 | F (2, 12) = 27.64 | p<0.0001 |
| Figure 2C | Drug on WIN effect | 7-40 | One-way ANOVA (Tukey) | CB1- vs Veh vs TG vs 2APB vs i11 | F (4, 104) = 9.818 | p<0.0001 |
| Figure 2D | Drug on WIN effect | 7-41 | One-way ANOVA (Tukey) | CB1- vs Veh vs TG vs 2APB vs i11 | F (4, 105) = 13.69 | p<0.0001 |
| Figure 2F | WIN on genotype/WIN vs Veh | 13-20 | Two-way ANOVA (Tukey) | Genotype vs WIN Interaction | F (2, 94) = 5.571 | 0.0052 |
| Figure 2G | Drug on WIN effect | 6-12 | One-way ANOVA (Tukey) | CB1- vs Veh vs MK | F (2, 25) = 4.995 | 0.0150 |
| Figure 2H | Drug on WIN effect | 7-12 | One-way ANOVA (Tukey) | CB1- vs Veh vs MK | F (2, 26) = 9.029 | 0.0011 |
| Figure 2I | Expression on WIN effect | 10-12 | Kruskal-Wallis test | Ctrl vs dnMCU vs MICU1-S124A vs MICU1-WT | | 0.0134 |
| Figure 3B | Infection on Cell colocalization (Hp) | 6-9 | Unpaired t-test | S100β vs NeuN | t=27.52, df=13 | p<0.0001 |
| Figure 3B | Infection on Cell colocalization (Cx) | 5 | Unpaired t-test | S100β vs NeuN | t=30.58, df=8 | p<0.0001 |
| Figure 3G | Time vs Treatment (THC) vs Genotype (WT) | 4 | Friedman test | THC treatment | X2=1, DF= 1 | 0.3170 |
| Figure 3G | Time vs Treatment (THC) vs Genotype (KO) | 4 | Friedman test | THC treatment | X2=1.DT=1 | 0.3170 |
| Figure 3H | Time vs Treatment (THC) vs Genotype (WT) | 4 | Friedman test | THC treatment | X2=4, DF= 1 | 0.0460 |
| Figure 3H | Time vs Treatment (THC) vs Genotype (KO) | 4 | Friedman test | THC treatment | X2=1.DT=1 | 0.3170 |
| Figure 3K | Treatment (THC) vs Genotype | 6-8 | Repeated two-way ANOVA (Sidak) | THC treatment vs genotype | F (1, 12) = 0.4211 | 0.5286 |
| Figure 3L | Treatment (THC) vs Genotype | 9 | Repeated two-way ANOVA (Sidak) | THC treatment vs genotype | F (1, 16) = 1.327 | 0.2663 |
| Figure 3M | Treatment (THC) vs Genotype | 6-8 | Repeated two-way ANOVA (Sidak) | THC treatment vs genotype | F (1, 12) = 6.138 | 0.0291 |
| Figure 3N | Treatment (THC) vs Genotype | 9 | Repeated two-way ANOVA (Sidak) | THC treatment vs genotype | F (1, 16) = 5.064 | 0.0388 |
| Figure 4C | DN22 genotype % potentiated synapses | 19-20 | Chi-square | Probability of potentiation | χ2=4.792, df=1 | 0.0286 |
| Figure 4E | ND on success rate (WT) | 19 | Two tailed paired t-test | Basal vs post ND | t=2.191, df=18 | 0.0419 |
| Figure 4E | ND on success rate (DN22-CB1-KI) | 20 | Two tailed paired t-test | Basal vs post ND | t=0.6578, df=19 | 0.5186 |
| Figure 5B | Stimulation vs Treatment (MCU-i11) | 6-7 | Repeated two-way ANOVA (Sidak) | Stimulation vs treatment | F (1, 11) = 0.3767 | 0.5519 |
| Figure 5B | Stimulation vs Treatment (MCU-i11) | 6-7 | Repeated two-way ANOVA (Sidak) | Stimulation | F (1, 11) = 5.765 | 0.0352 |
| Figure 5C | Stimulation vs Treatment (MCU-i11) | 6-7 | Repeated two-way ANOVA (Sidak) | Stimulation vs treatment | F (1, 11) = 5.003 | 0.0470 |
| Figure 5C | Stimulation vs Treatment (MCU-i11) | 6-7 | Repeated two-way ANOVA (Sidak) | Stimulation | F (1, 11) = 7.366 | 0.0201 |
| Figure 5D | Stimulation vs Treatment (MCU-i11) | 6-7 | Repeated two-way ANOVA (Sidak) | Stimulation vs treatment | F (1, 11) = 6.160 | 0.0305 |
| Figure 5D | Stimulation vs Treatment (MCU-i11) | 6-7 | Repeated two-way ANOVA (Sidak) | Stimulation | F (1, 11) = 6.957 | 0.0231 |
| Figure 5E | Stimulation vs Treatment (MCU-i11) | 6-7 | Repeated two-way ANOVA (Sidak) | Stimulation vs treatment | F (1, 11) = 2.938 | 0.1145 |
| Figure 5E | Stimulation vs Treatment (MCU-i11) | 6-7 | Repeated two-way ANOVA (Sidak) | Stimulation | F (1, 11) = 11.10 | 0.0067 |
| Figure 5F | MCU inh vs % potentiated synapses | 19-26 | Chi-square | Probability of potentiation | χ2=8.024, df=3 | 0.0455 |
| Figure 5H | ND on success rate (Ctrl) | 21 | Two tailed paired t-test | Basal vs post ND | t=2.698, df=20 | 0.0138 |
| Figure 5H | ND on success rate (i11) | 19 | Two tailed paired t-test | Basal vs post ND | t=1.069, df=18 | 0.2991 |
| Figure 5J | ND on success rate (GFP) | 26 | Two tailed paired t-test | Basal vs post ND | t=2.721, df=25 | 0.0117 |
| Figure 5J | ND on success rate (DN-MCU) | 26 | Two tailed paired t-test | Basal vs post ND | t=1.016, df=25 | 0.3194 |

**Table S1**

| Figure | Conditions | "n" | Analysis (post-hoc test) | Factor analyzed | | P value |
|---|---|---|---|---|---|---|
| Figure S1B | Transfection on mean intensity | 5-9 | Unpaired t-test | CB1 vs DN22-CB1 | t=1.185, df=12 | 0.2588 |
| Figure S1C | Transfection on colocaliztion | 2-14 | One-way ANOVA (Tukey) | Ctrl vs CB1 vs DN22-CB1 | F(2,22)=8.50 | 0.0018 |
| Figure S1E | Tranferction on distance | 9-13 | Repeated two-way ANOVA (Sidak) | Transfection vs distance | F (2, 40) = 8.472 | 0.0009 |
| Figure S2A | ATP on Genotype | 5 | One-way ANOVA (Tukey) | Ctrl vs CB1 vs DN22-CB1 | F (2, 12) = 0.2263 | 0.8008 |
| Figure S3B | Virus on % mitochondria | 10 | One-way ANOVA (Tukey) | Ctrl vs CB1 vs DN22-CB1 | F (2, 27) = 2,687 | 0.0863 |
| Figure S3C | Virus on perimeter | 9 | One-way ANOVA (Tukey) | Ctrl vs CB1 vs DN22-CB1 | F (2, 24) = 0,5566 | 0.5804 |
| Figure S3D | Virus on Ca_score | 10 | One-way ANOVA (Tukey) | Ctrl vs CB1 vs DN22-CB1 | F (2, 27) = 0,3466 | 0.7102 |
| Figure S5B | Treatment (THC) vs Genotype (duration) | 6-8 | Repeated two-way ANOVA (Sidak) | THC treatment vs genotype | F (1, 12) = 0.6751 | 0.4273 |
| Figure S5B | Treatment (THC) vs Genotype (frequency) | 6-8 | Repeated two-way ANOVA (Sidak) | THC treatment vs genotype | F (1, 12) = 2.124 | 0.1707 |
| Figure S5C | Treatment (THC) vs Genotype (duration) | 8-9 | Repeated two-way ANOVA (Sidak) | THC treatment vs genotype | F (1, 15) = 0.2281 | 0.6398 |
| Figure S5C | Treatment (THC) vs Genotype (frequency) | 9 | Repeated two-way ANOVA (Sidak) | THC treatment vs genotype | F (1, 16) = 0.3876 | 0.5423 |
| Figure S5D | Treatment (THC) vs Genotype (duration) | 6-8 | Repeated two-way ANOVA (Sidak) | THC treatment vs genotype | F (1, 12) = 3.764 | 0.0762 |
| Figure S5D | Treatment (THC) vs Genotype (frequency) | 6-8 | Repeated two-way ANOVA (Sidak) | THC treatment vs genotype | F (1, 12) = 0.9337 | 0.3530 |
| Figure S5E | Treatment (THC) vs Genotype (duration) | 9 | Repeated two-way ANOVA (Sidak) | THC treatment vs genotype | F (1, 16) = 2.780 | 0.1149 |
| Figure S5E | Treatment (THC) vs Genotype (frequency) | 9 | Repeated two-way ANOVA (Sidak) | THC treatment vs genotype | F (1, 16) = 0.0008515 | 0.9771 |
| Figure S6A | ND on EPSC amplitude (WT) | 19 | Two tailed paired t-test | Basal vs post ND | t=0.1790, df=18 | 0.8600 |
| Figure S6A | ND on EPSC amplitude (DN22-CB1-KI) | 20 | Two tailed paired t-test | Basal vs post ND | t=1.386, df=19 | 0.1819 |
| Figure S6B | Genotype on success rate | 19-20 | Two tailed unpaired t-test | WT vs DN22-CB1-KI | t=0.7689, df=37 | 0.4469 |
| Figure S6C | Genotype on success rate | 19-20 | Two tailed unpaired t-test | WT vs DN22-CB1-KI | t=0.2892, df=37 | 0.7740 |
| Figure S7C | Stimulation vs Treatment (MCU-i11) | 6-7 | Repeated two-way ANOVA (Sidak) | Stimulation vs treatment | F (1, 11) = 0.1862 | 0.6745 |
| Figure S7D | Stimulation vs Treatment (MCU-i11) | 6-7 | Repeated two-way ANOVA (Sidak) | Stimulation vs treatment | F (1, 11) = 0.009018 | 0.9261 |
| Figure S7E | Stimulation vs Treatment (MCU-i11) | 6-7 | Repeated two-way ANOVA (Sidak) | Stimulation vs treatment | F (1, 11) = 1.797 | 0.2071 |
| Figure S7F | Stimulation vs Treatment (MCU-i11) | 6-7 | Repeated two-way ANOVA (Sidak) | Stimulation vs treatment | F (1, 11) = 0.002175 | 0.9636 |
| Figure S8A | ND on EPSC amplitude (Ctrl) | 21 | Two tailed paired t-test | Basal vs post ND | t=0.7059, df=20 | 0.4884 |
| Figure S8A | ND on EPSC amplitude (i11) | 19 | Two tailed paired t-test | Basal vs post ND | t=0.2287, df=18 | 0.8217 |
| Figure S8A | ND on EPSC amplitude (GFP) | 26 | Two tailed paired t-test | Basal vs post ND | t=1.827, df=25 | 0.0796 |
| Figure S8A | ND on EPSC amplitude (DN-MCU) | 26 | Two tailed paired t-test | Basal vs post ND | t=1.081, df=25 | 0.2899 |
| Figure S8B | Drug on EPSC amplitude | 21-19 | Two tailed unpaired t-test | Ctrl vs i11 | t=0.05709, df=38 | 0.9548 |
| Figure S8B | Virus on EPSC amplitude | 26-26 | Two tailed unpaired t-test | GFP vs dnMCU | t=1.355, df=50 | 0.1816 |
| Figure S8C | Drug on success rate | 21-19 | Two tailed unpaired t-test | Ctrl vs i11 | t=2.196, df=38 | 0.0343 |
| Figure S8C | Virus on success rate | 26-26 | Two tailed unpaired t-test | GFP vs dnMCU | t=0.3541, df=50 | 0.7247 |

**Table S2**
